# The temporal organization of memory and emotion is reciprocally coupled

**DOI:** 10.64898/2026.03.12.711172

**Authors:** Mengsi Li, Brooke Schwartzman, Tori N. LeVier, Barry Giesbrecht, Regina C. Lapate

## Abstract

Emotional experiences unfold over time and can bias the appraisal and memory of subsequent events. Yet, it remains unclear whether and how the temporal organization of memory relates to the persistence of affect over time, or affective spillover—and conversely, how emotional-valence dynamics sculpts temporal memory. We developed a novel paradigm employing positive and negative emotional sequences and combined EEG and behavioral measures of temporal memory and affective spillover. Longer remembered temporal distance for emotional-sequence items was associated with reduced affective spillover onto later-shown neutral stimuli, whereas better order memory correlated with greater spillover. Alpha burst time during emotional-sequence encoding predicted longer remembered temporal distance and modulated spillover in a valence-dependent manner. Conversely, emotional-valence shifts expanded remembered temporal distance and enhanced order memory across sequences. These findings demonstrate reciprocal coupling between temporal memory organization and affect dynamics, revealing how memory structure shapes—and is shaped by—the persistence of emotion.

## Introduction

Emotions are dynamic, and often persist beyond the moments they arise, shaping how we perceive and remember the unfolding stream of daily life^1–6^. For instance, the frustration following the sudden discovery that your bike has been stolen can linger for hours and make a colleague’s otherwise neutral remark on a manuscript later that day seem harsh (**Fig. 1a**). This phenomenon—whereby emotional responses linger beyond the duration of their original triggers and bias the perception and appraisal of subsequent, unrelated experiences—is referred to as *affective spillover*^7–10^. Beyond shaping appraisal, recent work underscores that emotions provide a powerful contextual signal that structures memory, segmenting continuous experience into discrete yet temporally structured events, and binding related experiences together in memory^11,12^. However, it remains unclear whether the temporal organization of valenced emotional events in memory influences the extent to which emotional responses “spill over” to subsequent experiences, and conversely, how changes in valence across emotional events shape their temporal organization in memory. Here, we examined this reciprocal, dynamic interplay between emotional responding and temporal memory. We tested whether high-fidelity temporal memory coding for emotional events was associated with reduced affective spillover, and conversely, whether dynamic emotional-valence shifts sculpted the temporal organization of emotional events in memory, while using scalp-recorded electroencephalography (EEG) to identify the neural correlates of online temporal memory coding and affect dynamics.

**Fig. 1.**
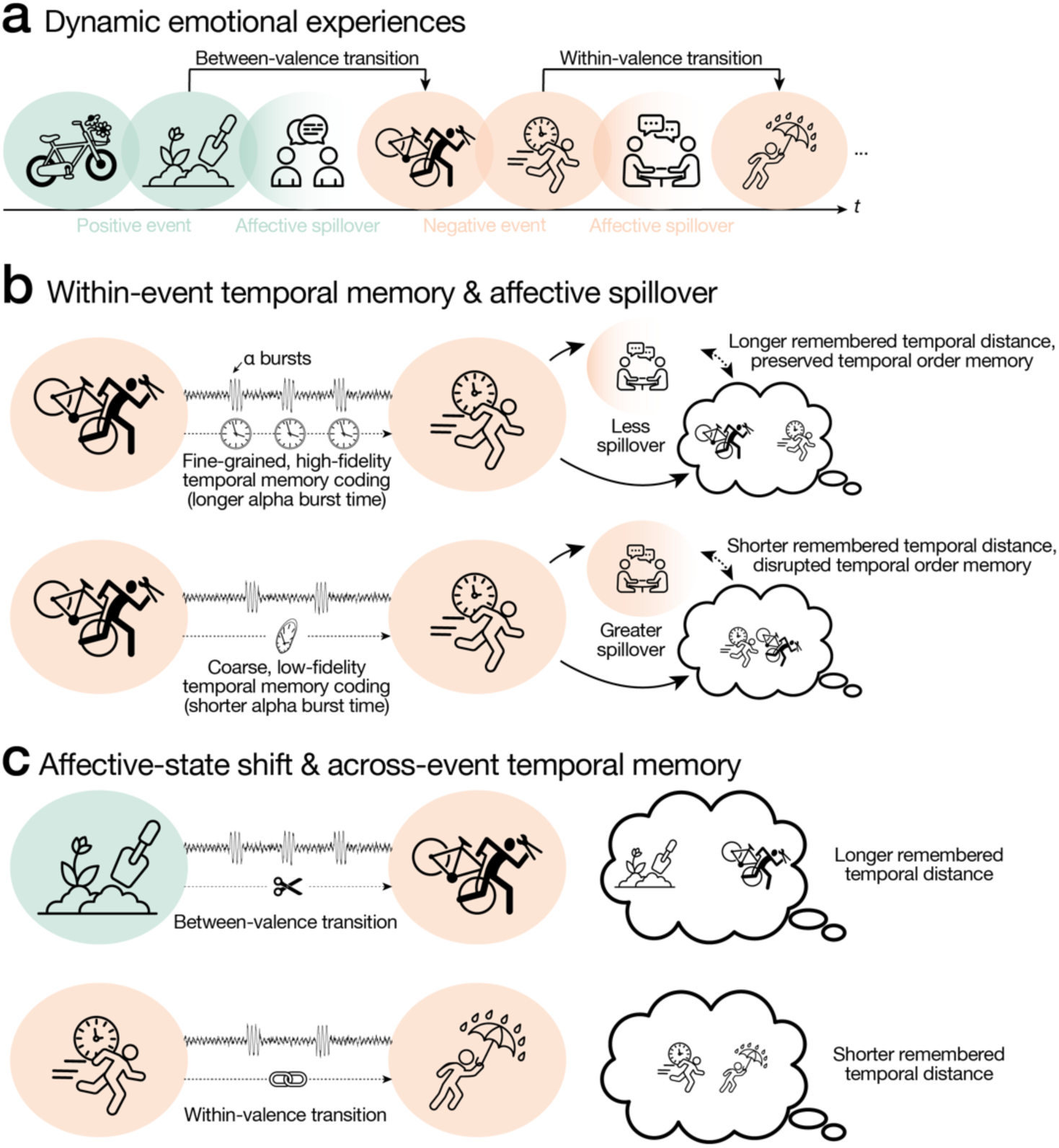
Conceptual framework of the hypotheses tested in this study. **a)** Everyday experience unfolds as a stream of positive and negative emotional events interleaved with neutral moments. We focused on two aspects of affect dynamics during emotional experiences: *affective spillover*, whereby emotional responses persist beyond a provocative event and bias appraisals of subsequent neutral experiences, and *affective-state shifts*, operationalized here as shifts in emotional-event valence. We tested the hypothesis that **b)** emotional events associated with fine-grained, high-fidelity temporal memory coding—indexed by greater remembered temporal distance and better order memory^13,21–25^—would also be associated with reduced subsequent affective spillover^13^. We further predicted that **c)** emotional-valence shifts (between-valence *vs.* within-valence transitions) would dilate remembered temporal distance *across* events^63,64^. We opted not to make directional predictions for temporal order memory given mixed prior findings^63–65^. Finally, we examined whether *alpha bursts* indexed online temporal memory coding (potentially reflecting internal state fluctuations), such that greater alpha burst time would predict expanded remembered temporal distance^66,67^ (**b** & **c**), and we also tested whether this putative neural signature of temporal memory coding **b)** was related to affective spillover and **c)** was modulated by emotional-valence shifts. Icons used in this figure were sourced from https://thenounproject.com.

Theoretical frameworks on emotional memory posit a trade-off between amygdala-dependent emotional processing and hippocampal-based contextual coding^11,13–20^, including temporal context coding such as temporal order^21–23^ and temporal distance^24,25^ across events. Consistent with this view, a growing empirical literature demonstrates emotion-driven distortions in temporal memory^26–32^ (but see also^33–35^; for reviews, see^11,13,14,36,37^). This trade-off raises the possibility that when temporal coding is impaired by emotional processing, emotional responses may become less anchored to their original temporal context and persist over time. Evidence obtained from individuals with mood and anxiety disorders aligns with this possibility, as they are often characterized by alterations in the persistence of emotional reactions as well as in temporal memory^13^. For instance, post-traumatic stress disorder (PTSD) and depression are characterized by the persistence—rather than the mere expression—of negative emotional responses over time (e.g., fear in PTSD and hopelessness in depression^1,4,38^). Furthermore, individuals with depression or PTSD often recall autobiographical memories in a temporally disorganized manner^38–40^. Likewise, individuals with higher dispositional negativity, a known risk factor for mood and anxiety disorders^41^, often recall negative events as lasting longer than they originally did^42^. Compromised structural integrity of the hippocampus is often observed in, and confers risk for, depression and PTSD^43–47^. Hippocampal lesions have also been linked to sustained negative affect following an emotionally provocative event^48^. Despite suggestive evidence from psychopathology and hippocampal damage, no previous study to our knowledge has directly tested the link between the fidelity of temporal memory coding and affective persistence. Here, we tested the novel hypothesis that fine-grained, high-fidelity temporal memory coding of emotional events would be associated with reduced affective spillover to subsequent experiences.

A nascent literature provides further evidence for the reciprocal influence of emotion on temporal memory, indicating that dynamic emotional states can shape the temporal organization of experiences in memory (for reviews, see^11,12^). Seminal work, originally conducted in non-emotional contexts, demonstrates that contextual shifts (such as moving to a different location) create *event boundaries* that structure memory (for reviews, see^49–52^), expanding remembered temporal distance,^25,53–55^ and often impairing order memory^53,56–58^ (but see^55,59^). Recent studies suggest that emotional stimuli at otherwise neutral event boundaries modulate these effects^60–62^, and that emotional-state transitions themselves induce boundary-like segmentation that dilates remembered temporal distance^63,64^. Yet, the extent to which emotion-induced boundary-like effects are driven by changes in emotional valence or arousal remains unclear because most prior work has contrasted transitions between negative and neutral states, which does not permit fully disentangling the effects of valence and arousal^64^. While McClay et al. (2023) employed positive and negative stimuli, which revealed boundary effects that were primarily driven by changes in subjective emotional valence rather than arousal, the effects they reported focused primarily on transitions *within* positive or negative states, as opposed to *between* positive and negative emotional valences^63^. Finally, while transitioning toward a more negative state—such as from neutral to negative^64^ or from less to more negative^63^—typically dilates remembered temporal distance, the nature of the impact of those transitions on temporal order memory remains unclear (with prior evidence for both enhancement^64,65^ and impairment^63^). Thus, examining dynamic transitions *between* positive and negative states—ubiquitous in daily life—while controlling for concurrent changes in arousal is essential for understanding how shifts in emotional-state valence influence temporal distance and order memory.

Bursting EEG alpha activity, recently implicated in the automatic tracking of time as experience unfolds (i.e., *episodic timing*^66–68^), may provide a neural correlate of online temporal memory coding. Early models proposed that the alpha rhythm functions as an internal pacemaker, oscillating at a steady rate and tracking the passage of metric time (much like a clock; the *alpha clock hypothesis*^69–72^). Emerging evidence, however, suggests that alpha activity is not *stationary*^73,74^, but instead occurs in transient bursts that may track internal “events” rather than metric time per se. Specifically, during awake rest, a larger proportion of time spent in *alpha bursts* (i.e., *alpha burst time*) predicts longer retrospective estimates of elapsed time, a finding interpreted as suggesting that alpha bursts reflect discrete “bouts of awareness” that might timestamp experience and ultimately support the temporal organization of events in memory^66–68^. Building on these findings, we tested whether alpha bursts tracked memory for temporal distance (and potentially for temporal order) in emotional events.

To address these aims, we developed a novel within-subjects paradigm employing dynamically changing negative and positive emotional events that allowed us to measure temporal memory for those events as well as the subsequent affective spillover they elicited. In the *Emotional Sequence Spillover Task*, participants viewed sequences (“*events*”) comprising four positive or negative images (“*items*”) interleaved with novel neutral faces. Following each sequence, participants rated the likeability of a novel neutral face, indexing affective spillover. We manipulated emotional-sequence transitions to create within-valence shifts as well as between-valence shifts across positive and negative states (**Fig. 2b**). After every 14 sequences, we assessed temporal distance and order memory for image pairs sampled from within and across events (see **Fig. 2c** and *Temporal memory task* in *Methods* for details). We recorded EEG throughout the task to investigate the role of alpha bursts during emotional-sequence encoding, providing a putative online neural correlate of subsequent temporal memory.

**Fig. 2.**
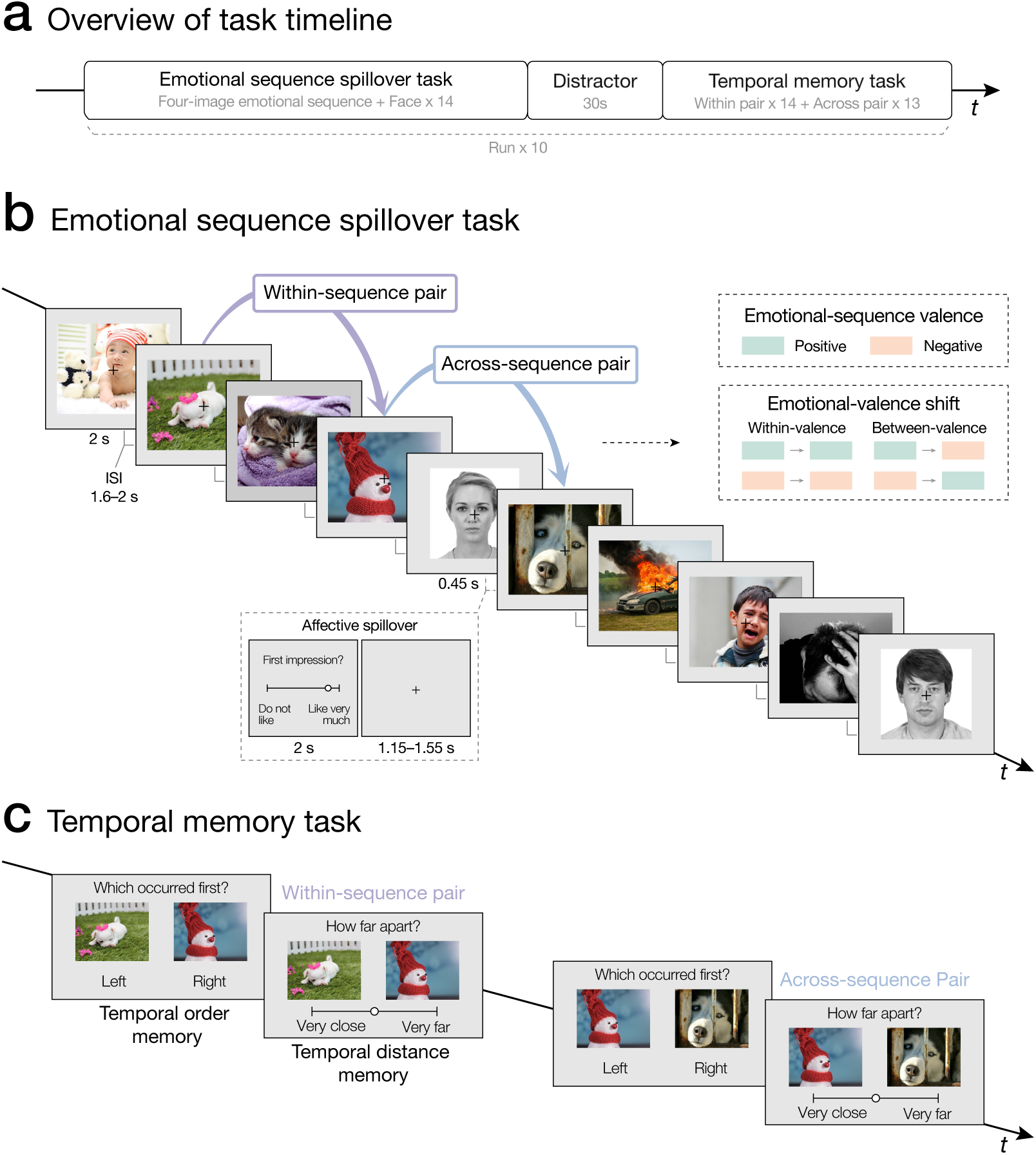
Schematic of the task design. **a)** Participants completed 10 runs of experimental tasks, each comprising a set of the *Emotional Sequence Spillover* and *Temporal Memory Tasks*, separated by a 30-s distractor interval. **b)** In each run of the *Emotional Sequence Spillover Task*, participants viewed 14 emotional sequences (“*events*”) consisting of four positive or negative images (“*items*”) interleaved with novel neutral faces. Following each sequence, participants rated the likeability of the neutral faces, which indexed affective spillover. We manipulated emotional-sequence transitions to create within-valence shifts as well as between-valence shifts across positive and negative states. Each emotional image was followed by a jittered inter-stimulus interval (1.6–2.0 s), and each face was followed by a 2-s rating period and a jittered inter-trial interval (1.15–1.55 s). **c)** After every 14 trials of the *Emotional Sequence Spillover Task* and a distractor, participants completed a *Temporal Memory Task* assessing memory for temporal order (“*Which occurred first?*”) and temporal distance (“*How far apart?*”) for image pairs sampled either within the same sequence or across adjacent sequences. Within- and across-sequence image pairs were matched on objective temporal distance. EEG was recorded throughout. The emotional images used in this figure are from the Open Affective Standardized Image Set (OASIS)^75^. The face images are from the Chicago Face Database (CFD)^76^.

We first tested whether emotional events associated with fine-grained, high-fidelity temporal memory coding—indexed by longer remembered temporal distance and better order memory for within-sequence image pairs^13,21–25^—were also associated with reduced affective spillover onto subsequent neutral faces^13^ (**Fig. 1b**). We then examined whether and how emotional-valence shifts shaped temporal memory *across* events. We predicted that between-valence emotional shifts would be associated with dilated remembered temporal distance across events, with additional dilation for positive-to-negative transitions^63,64^ (**Fig. 1c**). We opted not to make directional predictions for temporal order memory given mixed prior findings^63–65^. Finally, we tested whether longer alpha burst time during the encoding of emotional events was associated with expanded remembered distance within and across events^66,67^, as well as whether this putative neural index of temporal memory coding was sensitive to shifts in emotional valence and correlated with the magnitude of subsequent affective spillover (**Fig. 1b & c**).

## Results

### Emotional sequences produce affective spillover

We first examined whether emotional sequences produced affective spillover, as indexed by whether likeability ratings of novel neutral faces were biased by the emotional valence of the preceding emotional sequences. We found that emotional sequences robustly produced affective spillover: participants rated novel neutral faces following positive emotional sequences as more likeable than neutral faces following negative emotional sequences (**Fig. 3a**; *B* = 0.14 (*SE* = 0.04), 95% CI [0.06, 0.23], *F* = 10.68, *p* = 0.002; *M*_Positive_ = 2.52; *SE*_Positive_ = 0.03; *M*_Negative_ = 2.38; *SE*_Negative_ = 0.03). This result persisted after controlling for neutral-face gender and normative likeability ratings (obtained from an independent sample; N = 100; see *Primary analyses: Affective spillover* in *Methods*)—thereby underscoring the robustness of affective spillover effects (*B* = 0.14 (*SE* = 0.04), 95% CI [0.06, 0.23], *F* = 10.66, *p* = 0.002; for additional details, see *Affective spillover: Face characteristics* in *Supplementary Results*). This finding extends prior work demonstrating affective spillover from a single emotional image to temporally extended emotional sequences^7–9^.

**Fig. 3.**
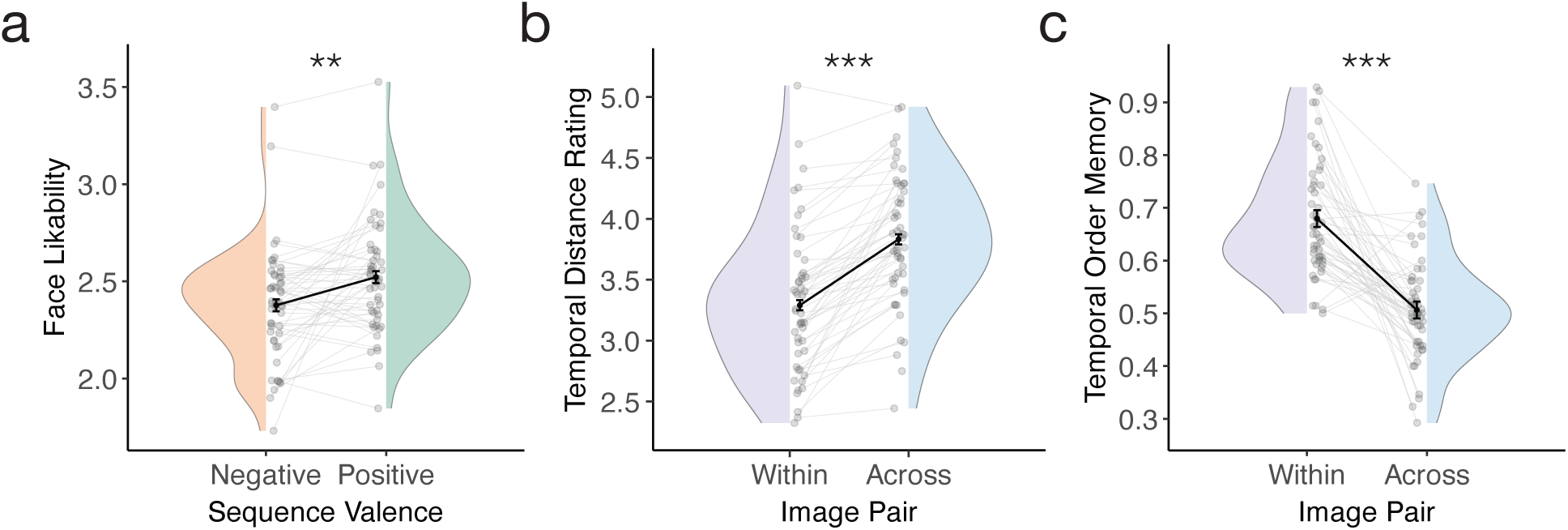
Affective spillover and event-boundary effects. **a)** Emotional sequence valence significantly predicted subsequent face likeability ratings, with faces following positive sequences rated as more likable than those following negative sequences—indicating robust affective spillover (*F* = 10.68, *p* = 0.002). **b)** Image pairs sampled *within* emotional sequences were remembered as closer in time than pairs sampled *across* sequences spanning a neutral face, demonstrating event-boundary effects on temporal distance memory (*F* = 86.39, *p* < 0.001). **c)** Temporal order memory was enhanced for within-relative to across-sequence pairs, further suggesting event-boundary effects on temporal memory (*χ²*(1) = 54.03, *p* < 0.001). Half-violin plots depict participant-level data. Light gray dots and connecting lines indicate individual participants’ datapoints across conditions, while black dots and connecting lines denote participant means. Error bars reflect ±1 SEM of the within-subject difference between conditions. \*\**p* < 0.01; \*\*\**p* < 0.001.

### Temporal memory is shaped by neutral stimuli embedded in emotional sequences

Next, we examined whether and how transitions between emotional sequences—marked by an intervening neutral face between sequences—modulated temporal distance and temporal order memory. Participants remembered image pairs sampled within emotional sequences as closer in time than pairs sampled across sequences—i.e., spanned a neutral face—despite matched objective temporal distance (**Fig. 3b**; *B* = −0.54 (*SE* = 0.06), 95% CI [−0.66, −0.43], *F* = 86.39, *p* < 0.001; *M*_Within_ = 3.29; *SE*_Within_ = 0.04; *M*_Across_ = 3.83; *SE*_Across_ = 0.04). Temporal order memory accuracy, which was reliably above chance (*M* = 0.60, *t*(50) = 10.40, 95% CI [0.58, 0.61], *p* < 0.001), was higher for within-sequence than across-sequence pairs (**Fig. 3c**; *B* = 0.78 (*SE* = 0.11), 95% CI [0.57, 0.99], OR = 2.19, 95% CI [1.78, 2.70], χ²(1) = 54.03, *p* < 0.001; *M*_Within_ = 0.68; *SE*_Within_ = 0.02; *M*_Across_ = 0.51; *SE*_Across_ = 0.02). Importantly, these emotional-sequence boundary effects on temporal distance and order memory remained significant even when comparing within-*vs.* across-sequence pairs of the same valence (distance memory: *B* = −0.29 (*SE* = 0.04), 95% CI [−0.38, −0.20], *F* = 37.90, *p* < 0.001; order memory: *B* = 0.90 (*SE* = 0.12), 95% CI [0.67, 1.14], OR = 2.47, 95% CI [1.96, 3.11], χ²(1) = 58.13, *p* < 0.001), and were comparable across positive and negative sequences (n.s. valence × image-pair type interaction; distance memory: *B* = 0.01 (*SE* = 0.06), 95% CI [−0.11, 0.13], *F* = 0.04, *p* = 0.85; order memory: *B* = −0.11 (*SE* = 0.09), 95% CI [−0.29, 0.06], OR = 0.89, 95% CI [0.75, 1.07], χ²(1) = 1.56, *p* = 0.21)—thereby indicating that these effects are partly driven by the presentation of novel neutral faces (and their ratings) between sequences (*see below for additional effects associated with emotional-valence shifts*). Together, these findings replicate canonical event-boundary effects on temporal memory^25,53–58^ and extend them by showing how affectively heterogeneous experiences shape the temporal organization of memory.

### Temporal memory coding of emotional events is associated with affective spillover

Having established robust affective spillover effects and temporal memory modulation induced by emotional-sequence transitions, we next addressed the novel question of whether emotional events associated with fine-grained, high-fidelity temporal memory coding were also associated with reduced subsequent affective spillover produced by them. Specifically, we tested whether remembered temporal distance and temporal order for within-sequence image pairs predicted the magnitude of affective spillover from those sequences onto later-presented novel neutral faces (i.e., *trialwise;* computed as the valence-congruent deviation of in-task ratings from normative baseline ratings; for details, see *Primary analyses: Affective spillover* in *Methods*).

Consistent with our hypothesis, emotional sequences remembered as having longer temporal distance between items produced reduced affective spillover, as indexed by reduced emotion-driven ratings of subsequently-presented novel neutral faces (**Fig. 4a**; *B* = −0.01 (*SE* = 0.005), 95% CI [−0.02, −0.001], *F* = 4.88, *p* = 0.03). Interestingly, contrary to our prediction, emotional sequences associated with higher temporal order memory produced larger, rather than smaller, affective spillover (**Fig. 4b**; *B* = 0.03 (*SE* = 0.02), 95% CI [0.002, 0.07], *F* = 4.30, *p* = 0.04). These associations were comparable across positive and negative sequences (n.s. valence × temporal memory interaction; distance memory: *B* = 0.005 (*SE* = 0.01), 95% CI [−0.01, 0.02], *F* = 0.23, *p* = 0.63; order memory: *B* = 0.02 (*SE* = 0.03), 95% CI [−0.04, 0.09], *F* = 0.57, *p* = 0.45), indicating that the relationship between temporal memory and affective spillover was independent of emotional valence; these associations were also independent of arousal levels or the valence × arousal interaction (*p*s > 0.1; for details, see *Within-sequence temporal memory and affective spillover: Arousal effects in Supplementary Results*). Together, shorter remembered temporal distance and better temporal order memory were associated with greater affective spillover across contexts.

**Fig. 4.**
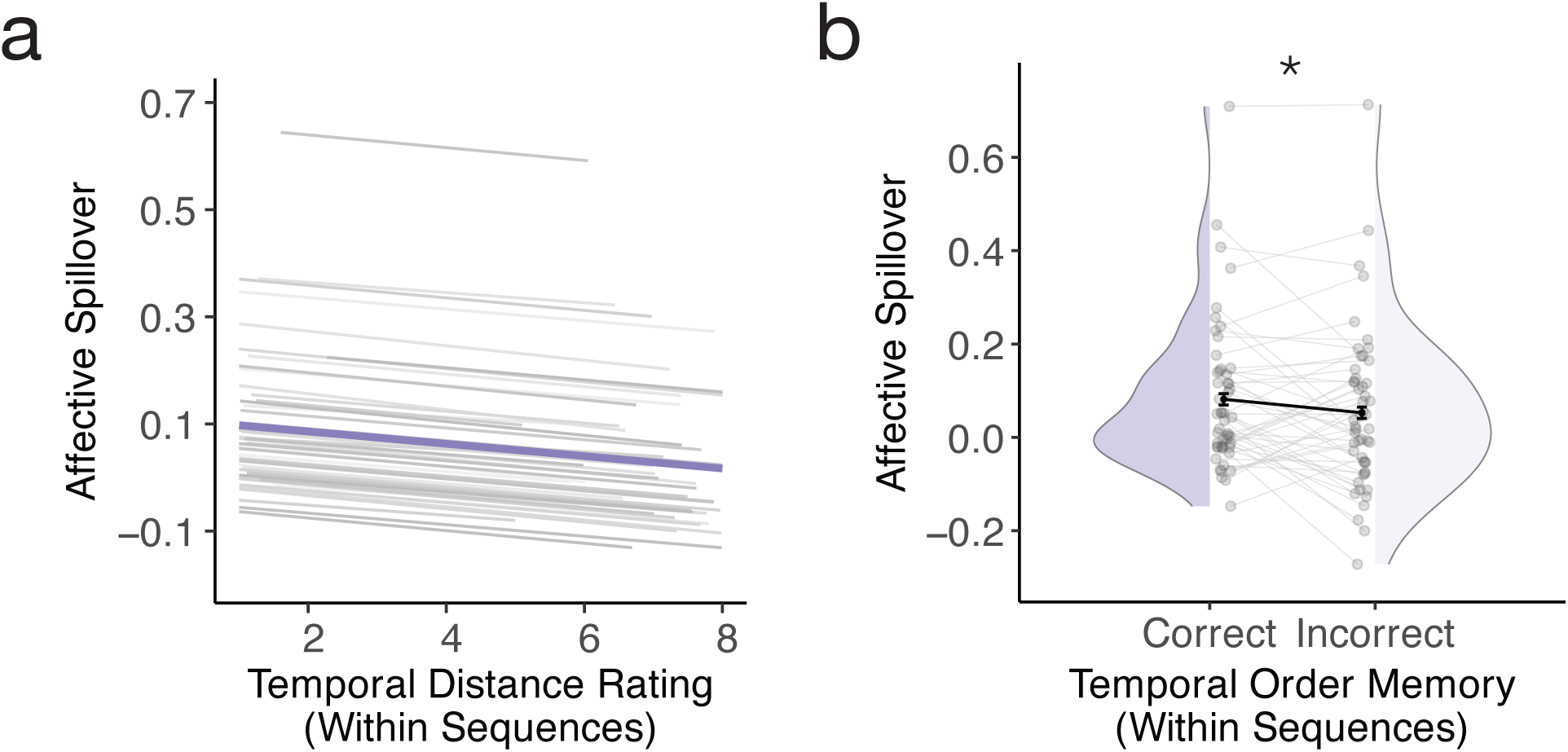
Temporal memory coding of emotional sequences is associated with affective spillover. Emotional sequences associated with **a)** shorter remembered temporal distance and **b)** correct order memory judgements produced greater affective spillover (distance memory: *B* = *−*0.01, *p* = 0.03; order memory: *F* = 4.30, *p* = 0.04). \**p* < 0.05.

### Alpha bursts track online temporal memory coding and subsequent affective spillover

#### Greater alpha burst time during emotional sequences correlates with longer remembered distance

Building upon recent evidence suggesting that alpha bursts at rest contribute to episodic timing^66,67^, we examined whether they might likewise support temporal memory coding during episodic emotional events. Alpha burst time was quantified as the proportion of time during emotional image viewing within a sequence in which alpha cycles were classified as robust oscillatory bursts^73^ (*trialwise* estimates averaged over an a-priori posterior cluster^66,67,77–79^ and baseline-corrected to ISI burst time; see **Fig. 5a & b** and *Methods: Alpha burst detection*).

**Fig. 5.**
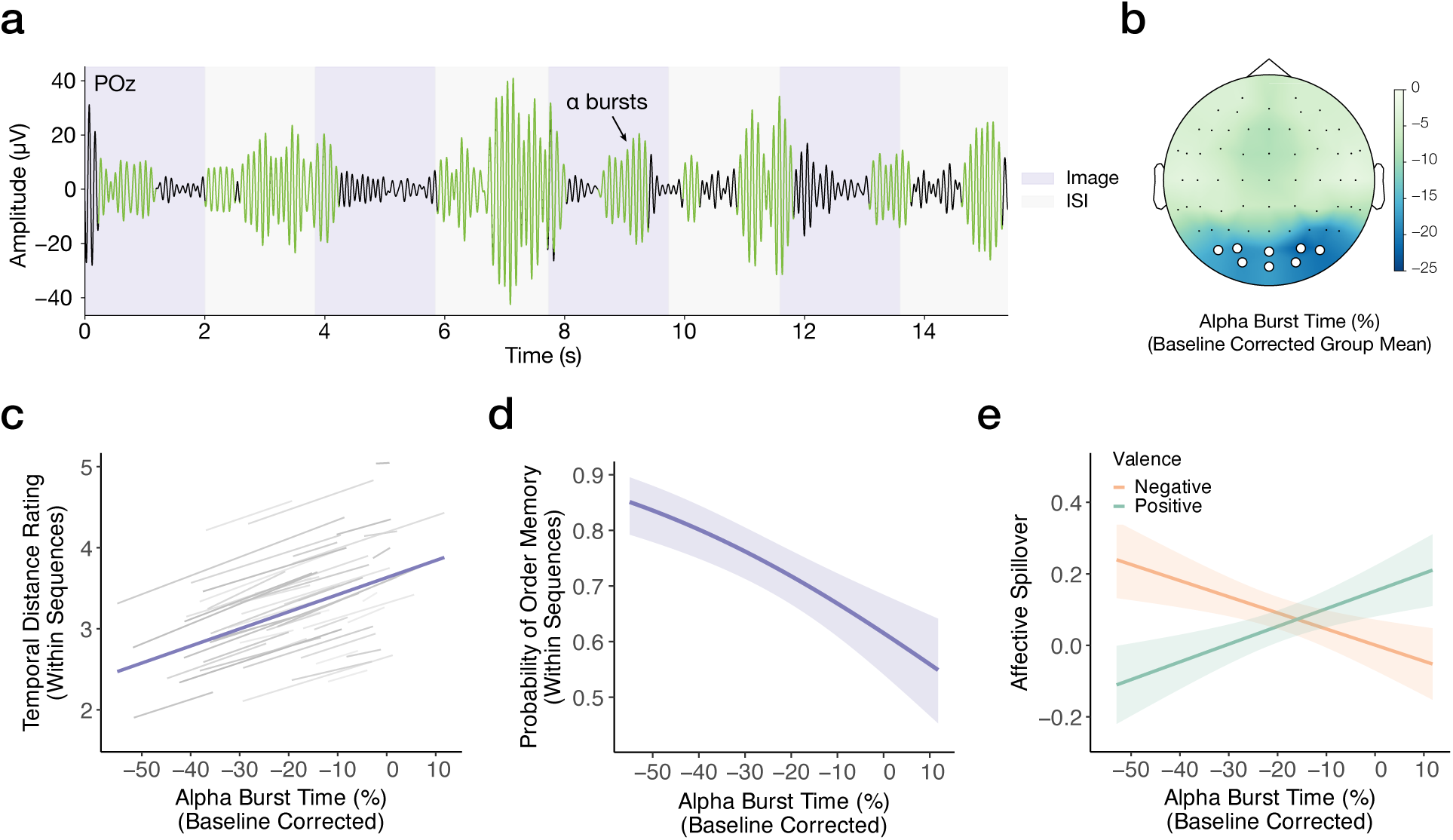
Alpha bursts track online temporal memory coding and subsequent affective spillover. **a)** *Trialwise* continuous EEG data—from each sequence onset through the post-sequence interval— were analyzed using the cycle-by-cycle approach^73^ to detect nonstationary alpha (8–12 Hz) bursts. The example trace illustrates alpha burst detection in an exemplar trial at POz. For each electrode, we computed *alpha burst time* as the proportion of time during which alpha activities were classified as oscillatory bursts, separately for image-presentation and ISI epochs. Stimulus-induced alpha burst time was then averaged across a predefined cluster of parieto-occipital electrodes and baseline-corrected by subtracting trial-averaged ISI burst time. **b)** The topomap shows the group-averaged distribution of baseline-corrected alpha burst time. We found that **c)** longer alpha burst time during emotional sequence encoding predicted longer remembered temporal distance for image pairs within the sequences (*B* = 2.11, *p*_FDR_ < 0.001). **d)** Longer alpha burst time was also associated with poorer within-sequence temporal order memory (*χ²*(1) = 23.31, *p*_FDR_ < 0.001). **e)** Finally, alpha burst time interacted with sequence valence in predicting affective spillover (*F* = 44.46, *p*_FDR_ < 0.001): longer burst time during negative sequences was associated with reduced spillover (*B* = *−*0.45, *p* = 0.003), whereas longer burst time during positive sequences predicted greater spillover (*B* = 0.50, *p* = 0.001).

We found that longer alpha burst time during emotional sequence encoding predicted longer remembered temporal distance for image pairs within the sequences (**Fig. 5c**; *B* = 2.11 (*SE* = 0.34), 95% CI [1.45, 2.78], *F* = 38.83, *p*_FDR_ < 0.001). Longer alpha burst time was also associated with poorer within-sequence temporal order memory (**Fig. 5d**; *B* = −2.31 (*SE* = 0.48), 95% CI [−3.25, −1.37], OR = 0.10, 95% CI [0.04, 0.25], χ²(1) = 23.31, *p*_FDR_ < 0.001). Importantly, these associations were specific to burst time obtained during emotional image viewing (not observed during the ISIs; distance memory: *B* = −0.26 (*SE* = 0.20), 95% CI [−0.64, 0.13], *F* = 1.66, *p*_FDR_ = 0.20; order memory: *B* = −0.57 (*SE* = 0.28), 95% CI [−1.13, −0.02], OR = 0.56, 95% CI [0.32, 0.98], χ²(1) = 4.14, *p*_FDR_ = 0.06). Furthermore, alpha burst time continued to significantly predict remembered temporal distance even after controlling for alpha power at the center frequency (i.e., alpha center power^80^; see *Methods: Alpha center power estimation*) (*B* = 1.80 (*SE* = 0.52), 95% CI [0.78, 2.82], *F* = 11.98, *p* = 0.0006)—suggesting the specificity of this association to nonstationary alpha rather than sustained oscillatory alpha activity. Interestingly, temporal order memory was instead most reliably associated with reduced alpha center power (i.e., greater event-related desynchronization, rather than alpha burst time; for details, see *Within-sequence temporal memory and affective spillover: Alpha center power* in *Supplementary Results*). Alpha center frequency, as well as burst time in the beta frequency band (13–30 Hz), was not significantly associated with either temporal memory measure, supporting the specificity of these effects to alpha-band bursting activity (*p*s > 0.05; for details, see *Within-sequence temporal memory: Specificity of alpha-burst effects* in *Supplementary Results*). Together, these findings suggest that alpha bursts support the tracking of episodic timing^66–68^ during emotional-event processing and may shape how emotional experiences are temporally structured in memory.

#### Greater alpha burst time during negative emotional sequences predicts reduced affective spillover

Next, we examined whether alpha burst time was associated with affective spillover and tested whether this association differed by sequence valence. We observed an interaction between alpha burst time and valence in predicting affective spillover, whereby longer alpha burst time during negative emotional sequences correlated with reduced affective spillover following those sequences (*B* = −0.45 (*SE* = 0.15), 95% CI [−0.75, −0.16], *t* = −3.00, *p* = 0.003). In contrast, longer alpha burst time during positive-sequence processing was associated with larger spillover following those sequences (*B* = 0.50 (*SE* = 0.15), 95% CI [0.20, 0.80], *t* = 3.26, *p* = 0.001; alpha burst time × valence: *B* = 0.95 (*SE* = 0.14), 95% CI [0.67, 1.22], *F* = 44.46, *p*_FDR_ < 0.001; **Fig. 5e**). Note, however, that this interaction was not predicted a priori, and therefore should be interpreted with caution. In addition, alpha burst time was not itself modulated by valence or arousal (valence: *B* = 0.002 (*SE* = 0.001), 95% CI [−0.0002, 0.005], *F* = 3.19, *p*_FDR_ = 0.10; arousal: *B* = −0.002 (*SE* = 0.002), 95% CI [−0.005, 0.0008], *F* = 2.14, *p*_FDR_ = 0.15), suggesting that it captured a valence-general temporal memory-coding process that modulates the persistence of affect in a valence-dependent manner. Alpha center power analyses revealed a similar interaction with valence in predicting affective spillover—with greater alpha power (i.e., weaker alpha desynchronization) predicting less spillover following negative sequences—as well as a positive association between sequence arousal and alpha desynchronization; for details, see *Within-sequence temporal memory and affective spillover: Alpha center power* in *Supplementary Results.* Collectively, these results suggest that alpha bursts may provide a proxy for a shared mechanism at the intersection of temporal memory coding and affective spillover.

#### Emotional-valence transitions shape temporal memory

Finally, we turned to the reciprocal influence of affect dynamics on temporal memory by testing whether shifts in emotional valence *across* sequences modulated temporal distance and order memory. We found that emotional-valence shifts dilated temporal distance memory (first *vs.* second emotional-sequence valence interaction; *F* = 184.08, *p* < 0.001; **Fig. 6a**), such that image pairs spanning an emotional-valence shift (positive-to-negative or negative-to-positive; *M* = 4.05, *SE* = 0.04) were remembered as farther apart than pairs without emotional-valence shifts (positive-to-positive or negative-to-negative; *M* = 3.58, *SE* = 0.04; *B* = 0.47 (*SE* = 0.03), 95% CI [0.40, 0.53], *t* = 13.56, *p* < 0.001). The direction of emotional-valence shift (positive-to-negative *vs.* negative-to-positive) did not differentially modulate these emotional-boundary effects on temporal distance (*B* = −0.06 (*SE* = 0.05), 95% CI [−0.15, 0.03], *t* = −1.25, *p* = 0.21). In sum, changes in emotional valence across sequences amplified event-boundary effects on temporal distance memory.

**Fig. 6.**
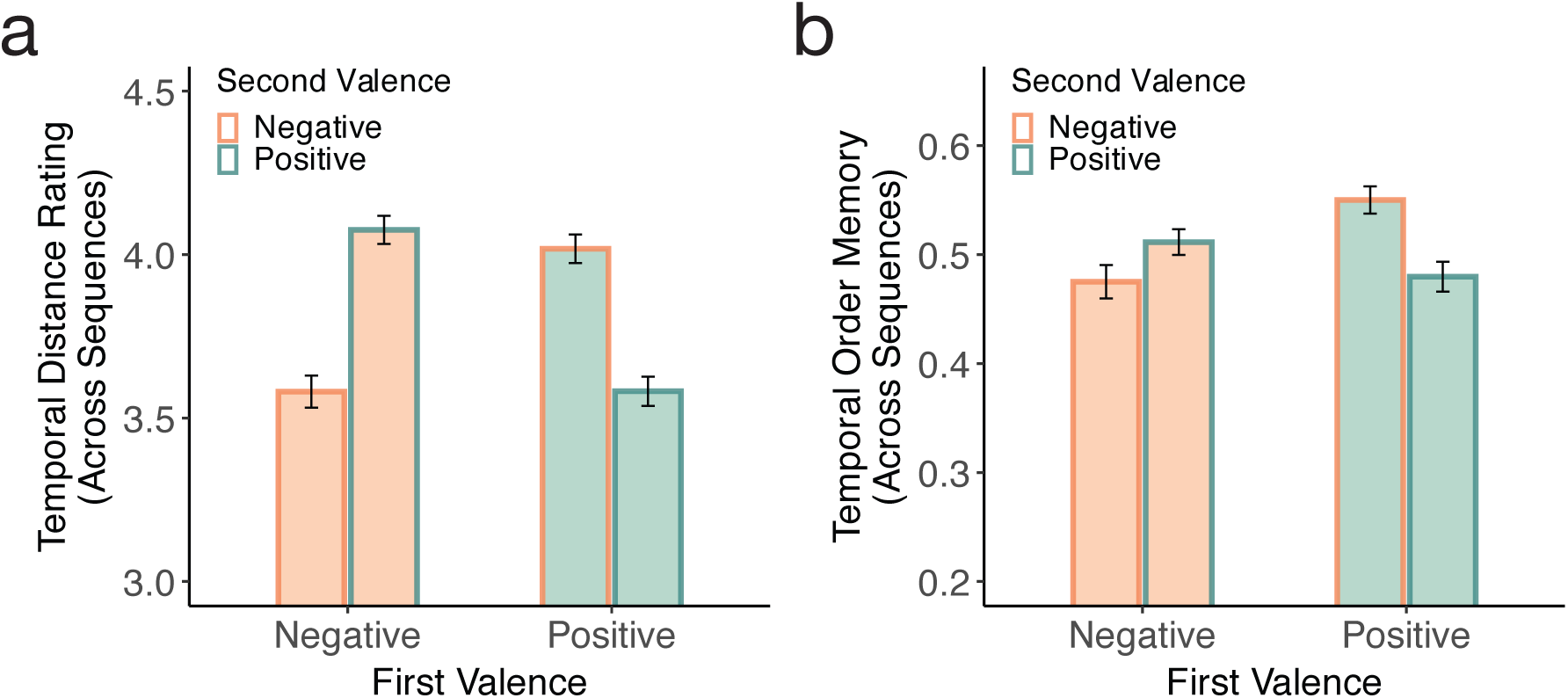
Emotional valence shifts differently shape temporal distance and order memory across event boundaries. **a)** Remembered temporal distance for across-sequence image pairs was modulated by an interaction between the valence of each sequence (first valence) and the valence of the subsequent sequence (second valence) (*F* = 184.08, *p* < 0.001). A comparison of pairs spanning a valence shift (positive-to-negative and negative-to-positive) with pairs that did not (positive-to-positive and negative-to-negative) revealed that emotional-valence shifts dilated remembered temporal distance (*t* = 13.56, *p* < 0.001). The direction of the emotional valence shift (positive-to-negative *vs.* negative-to-positive) did not additionally modulate this effect (*t* = *−*1.25, *p* = 0.21). **b)** Similarly, temporal order memory for across-sequence pairs was modulated by an interaction between the valence of the current sequence and the valence of the following sequence (*χ²*(1) = 19.32, *p* < 0.001), whereby emotional-sequence transitions spanning an emotional-valence shift were associated with better temporal order memory than boundaries without a valence shift (*z* = 4.40, *p* < 0.001). The direction of the emotional valence shifts additionally modulated the impact of emotional-valence transition on temporal order memory, with higher temporal order memory accuracy observed for positive-to-negative (compared to negative-to-positive) transitions (*z* = 2.35, *p* = 0.02). Error bars reflect ±1 SEM of the within-subject difference between conditions.

Emotional-valence transitions also robustly modulated temporal order memory (first *vs.* second emotional-sequence valence interaction; χ²(1) = 19.32, *p* < 0.001; **Fig. 6b**), such that image pairs spanning an emotional-valence shift (*M* = 0.53, *SE* = 0.01) were associated with better temporal order memory (compared to sequence transitions without a valence shift *M* = 0.48, *SE* = 0.01; *B* = 0.22 (*SE* = 0.05), 95% CI [0.12, 0.32], OR = 1.25, 95% CI [1.13, 1.37], *z* = 4.40, *p* < 0.001).

Critically, we found that the *direction* of emotional-sequence shift (positive-to-negative *vs.* negative-to-positive) modulated the influence of emotional-state changes on temporal order memory, such that *positive-to-negative* transitions (*M* = 0.55, *SE* = 0.01) produced higher temporal order memory accuracy compared to negative-to-positive ones (*M* = 0.51, *SE* = 0.01; *B* = 0.16 (*SE* = 0.07), 95% CI [0.03, 0.29], OR = 1.17, 95% CI [1.03, 1.34], *z* = 2.35, *p* = 0.02). Signal detection analyses revealed that this temporal order-memory advantage for positive-to-negative transitions was driven by enhanced order sensitivity for those positive-to-negative pairs (*d*′: *M* = 0.16, *t*(50) = 2.07, 95% CI [0.005, 0.32], *p* = 0.04) as well as by a response bias across subjects toward judging negative images as having occurred later (C: *M* = −0.05, *t*(50) = −2.66, 95% CI [−0.09, −0.01], *p* = 0.01). Together, these results show that emotional-valence transitions differentially modulate temporal memory: amplifying event-boundary effects on temporal distance memory, but attenuating boundary effects on temporal order memory, particularly for positive-to-negative shifts.

### Control analyses

Control analyses indicated that the associations between temporal memory and affective spillover, as well as the effects of emotional-valence transitions on temporal memory, remained robust after accounting for additional factors that can potentially influence temporal memory, including arousal^11,14,36^, semantic similarity^32,81–84^, and item exposure history during temporal memory testing. (See *Supplementary Results* for details, including analyses examining the independent contributions of these control factors as well as of alpha burst time elicited by images across sequences to temporal memory.)

## Discussion

Time provides a fundamental organizing dimension for both emotion and memory, along which emotional responses dynamically unfold and episodic events are structured. Despite this shared temporal scaffold, how the dynamics of affect and the temporal organization of emotional experiences in memory shape one another remains unclear. Here, we addressed this question using a novel paradigm in which positive and negative emotional sequences were interleaved with novel neutral faces, which served as a probe for the persistence or spillover of affect over time. We observed robust affective spillover from emotional sequences onto novel neutral faces, indicating that emotional states elicited during dynamic emotional-event processing persisted beyond their original context to shape the appraisal of later-presented, otherwise neutral stimuli^7–10^. Moreover, we replicated and extended canonical event-boundary effects on temporal distance and order memory, whereby emotional items separated by an intervening neutral face were remembered as farther apart in time and with lower temporal order memory accuracy than items encoded within the same emotional event^25,53–58^ (for reviews, see^11,12^). Critically, emotional sequences characterized by larger remembered temporal distance and lower temporal order memory were associated with reduced affective spillover. At the neural level, alpha burst time during emotional-event processing tracked trial-by-trial variation in temporal distance memory and was associated with affective spillover in a valence-dependent manner—with longer burst time predicting reduced spillover from negative emotional sequences. Complementing the association between temporal memory coding and affective persistence, shifts in emotional-state valence sculpted temporal memory, dilating remembered temporal distance and enhancing temporal order memory across events. Together, these findings provide converging behavioral and neural evidence that temporal memory coding and affect dynamics are closely and bidirectionally intertwined, and suggest that the temporal structuring of emotional experiences in memory may support the context-sensitive regulation of negative emotional responses over time.

We hypothesized that high-fidelity temporal memory coding—putatively reflected in longer remembered temporal distance and higher temporal order memory—would be associated with reduced affective spillover^13^. Consistently, we found that emotional sequences characterized by longer remembered temporal distance were associated with reduced spillover onto appraisals of novel neutral faces, suggesting that greater contextual separation between emotional items may help constrain emotional responses to their originating context. However, contrary to our hypothesis, better temporal order memory for items within a sequence was associated with greater (rather than less) spillover. Memory for temporal order and temporal distance likely reflects partially dissociable aspects of temporal memory organization, with potentially distinct implications for how emotion and temporal memory interact. Remembered temporal distance has been shown to reliably reflect the degree of contextual similarity between experiences, such that events encoded in similar contexts and/or associated with similar hippocampal activity patterns over time^24,25^ are often remembered as closer in time. Temporal order memory, on the other hand, is thought to be supported by mechanisms that vary as a function of the degree of contextual similarity between items^85–87^. For instance, when contextual similarity is high, order judgements may be primarily supported via associative or inter-item binding between items, whereby retrieval of one item cues its successors through local relational links^86,88^. In temporal memory experiments including the present task, where associative encoding across successive items is often encouraged, narrative construction through semantic and/or causal links may jointly facilitate greater contextual similarity and higher inter-item associative binding, which may in turn give rise to shorter temporal distance judgements^32,89^ and higher temporal order memory^59,90^ for within-event items. Consistently, temporal order memory is often disrupted for emotional items when they are encoded individually^29,30^, but maintained or enhanced when inter-item binding is supported through narrative encoding^34,35^ or whenever emotional experiences contain inherent narrative structure^33,91^. Collectively, our findings suggest that when distinct emotional items are represented as a more cohesively integrated and temporally organized event—as reflected in higher temporal order memory and shorter remembered temporal distance—emotional responses may be particularly prone to persisting beyond the emotional episode and biasing subsequent information processing.

At the neural level, our findings suggest that alpha bursts may index internal context fluctuations associated with the differentiation of experiences over time and support online temporal memory coding during emotional-event processing. Specifically, we found that longer single-trial alpha burst time during emotional-sequence processing predicted longer remembered temporal distance for items within those sequences. This finding converges with recent evidence that longer alpha burst time during quiet wakefulness predicts longer single-trial retrospective estimates of elapsed time^66,67^ and provides support for emerging accounts proposing that alpha bursts, which capture the nonstationary nature of alpha activity, reflect discrete internal states that timestamp endogenous events. This process, termed episodic timing, is thought to support the temporal organization of memory^66–68^. Collectively, these findings are also compatible with contemporary frameworks and empirical evidence indicating that gradually drifting neural population states intrinsically reflect time and may provide a temporal tag for events in memory^13,21,22,24,25,68,92–95^. In sum, alpha bursts measured with EEG may reveal a temporally resolved neural signature of internal context dynamics that structure episodic experience in memory.

Beyond temporal distance memory, alpha burst time also predicted affective spillover in a valence-specific manner: longer alpha burst time during negative events was associated with reduced spillover from negative (but not positive) emotional sequences. This finding suggests that alpha burst dynamics may reflect the contextual differentiation of emotional experiences in memory in a way that recapitulates the observed association between remembered temporal distance and reduced spillover: when internal context evolves more distinctly, as reflected in greater alpha burst time, negative emotional experiences may become more temporally segregated, which may help reduce the persistence and/or spillover of negative emotional responses to unrelated stimuli. Interestingly, longer alpha burst time was associated with greater spillover from positive emotional events, suggesting that the consequences of internal context dynamics for affective persistence may differ as a function of valence. It is possible that positive affect elicited by successive positive images promoted broader attentional focus^96–98^ and higher associative processing^99–101^ relative to the negative images, potentially facilitating the relational linkage of emotional experiences with subsequent stimuli despite ongoing internal contextual fluctuations. Note, however, that the differential association between alpha burst time and affective spillover by valence was not predicted, and thus should be replicated in future studies.

Of note, longer alpha burst time correlated with worse temporal order memory accuracy in this study. However, greater alpha center power (i.e., weaker event-related desynchronization) was likewise associated with worse temporal order memory accuracy—and when both measures were modeled jointly, only alpha power explained variance in temporal order memory accuracy, whereas alpha burst time uniquely predicted remembered temporal distance. Alpha desynchronization was further associated with higher normative sequence arousal, consistent with prior work linking alpha desynchronization to increased cortical excitability (e.g.^102–104^) and emotion-driven increases in attentional engagement (e.g.^105–108^). Together, these findings suggest that alpha burst dynamics and alpha power reflect partially dissociable mechanisms supporting the temporal organization of emotional experiences in memory, with alpha bursts more closely tracking internal context dynamics associated with remembered temporal distance, beyond the more generic processes related to attention and arousal^105–108^ typically reflected in alpha power.

We found that emotional-valence shifts across sequences dilated remembered temporal distance—even while controlling for changes in emotional arousal. Emerging evidence suggests that changes in emotional states drive event segmentation^63,64^ (for reviews, see^11,12^), akin to event-boundary effects induced by changes in external context^25,53,54,56,58^ (for reviews, see^49–52^). However, the relative contributions of emotional valence and arousal to these boundary-like effects remained unclear, as prior work has often confounded them by primarily contrasting negative and neutral stimuli (e.g.^64,65^; for an exception, see^63^; for reviews, see^11,12^). Together with prior work^63–65^, our findings suggest that emotional states may serve as a salient contextual signal that shapes temporal memory organization, consistent with current computational^16,81,109^ and theoretical models^12,14^. Our findings also showed that shifts in emotional valence sculpted remembered temporal distance above and beyond changes in arousal. Nonetheless, whether the temporal dilation effects associated with emotional-valence shifts reflect mechanisms specific to emotional valence *per se* or more general change-detection mechanisms triggered by changes in emotional context remains to be determined by future studies directly comparing temporal memory changes produced by emotional and non-emotional contextual transitions.

In contrast to classic event-segmentation findings, emotional-valence shifts across sequences *enhanced* (rather than impaired) temporal order memory. Whereas contextual changes induced by perceptual shifts (such as changes in a border color) are often associated with impaired order memory^53,56^, recent work suggests that order memory can be enhanced for items encoded across (*vs.* within) boundaries^55,59^ whenever those items provide diagnostic information of their original encoding contexts that scaffold temporal judgements^55^ (e.g. animals in zoos *vs.* artwork in galleries). In the present study, items within an emotional sequence shared a common valence, rendering valence a salient contextual cue bound to emotional items, which may have facilitated contextual retrieval and associative chaining processes^86,88^ supporting order memory judgements for items sampled across sequences. In addition, we observed greater temporal order memory for positive-to-negative than negative-to-positive transitions, driven by both greater sensitivity to negative events occurring later and a bias to judge negative items as having occurred more recently.

The following limitations of the present work warrant future investigation. First, the present findings underscored the interactions between temporal memory and emotional persistence as well as the role of emotional-valence transitions in sculpting temporal memory— while controlling for arousal using normative ratings of emotional images. However, given the critical role of arousal in memory^19,110^, and previous work implicating arousal-related event-boundary effects on temporal distance and order memory^53,60^, future work should include a neutral condition alongside positive and negative conditions while acquiring psychophysiological measures independently indexing valence and arousal, which will permit fully dissociating the unique and possibly interactive^60,63^ contributions of valence and arousal to temporal memory following emotional-state shifts. Including a neutral condition will also permit testing whether differences in temporal memory and alpha burst dynamics between emotional and neutral conditions predict affective spillover beyond those measures in the emotional condition alone.

Second, we observed dissociable effects of emotional valence on temporal distance and order memory across events, and these two facets of temporal memory showed distinct associations with subsequent affective spillover. Prior work suggests that the relationship between temporal distance and order memory itself depends on the strength of contextual similarity as well as inter-item associative binding during encoding^85,87^. Therefore, future work should systematically vary semantic and/or causal coherence between items to examine how contextual and associative coding shape temporal memory organization of emotional events and their downstream emotional consequences. Third, instructions to link emotional images into narratives—intended to facilitate temporal memory coding—may have influenced the degree to which temporal order memory depends on contextual differentiation vs. associative binding^85–87^, and in turn, shaped its relationship with subsequent affective spillover. Future studies should examine whether the association between temporal order memory and affective spillover observed in our study holds without instructions promoting inter-item binding via narratives. Finally, precisely how temporal coding and memory contribute to the regulation of affective responses likely varies across timescales, given ample evidence that memory for emotional content and contextual details follows distinct consolidation trajectories^111–113^. Thus, future work examining the interaction of emotion and temporal memory at both short and long timescales— ideally incorporating self-relevant, ecologically valid stimuli that elicit sustained emotional responses^91^—will be essential to fully determine the dynamics of these processes relevant for adaptive behavior and how they confer resilience or vulnerability to psychopathology.

In conclusion, our results provide converging neural and behavioral evidence revealing the dynamics of reciprocal interactions between temporal memory and emotion, whereby the temporal organization of emotional experiences in memory is linked to the persistence of emotion across contexts and shifts in emotional state reciprocally sculpt the temporal structure of those experiences in memory.

## Methods

### Participants

Fifty-two healthy adults participated in the current experiment. One participant was excluded from all analyses due to excessive EEG artifacts, resulting in a final sample of N = 51 (age range = 18–27, *M* = 19.31; *SD* = 1.68; 35 females). The sample size was not preregistered. Nevertheless, our sample size is substantially larger than typical EEG studies on related topics (affective spillover: N = 25^114^; emotion: N = 17–45^107,108^; temporal memory: N = 34^115^). It is also larger than or comparable to recent studies on alpha bursts and episodic timing (N = 26–57^66,67^).

In addition, we conducted a simulation-based power analysis to determine the sample size needed to achieve at least 80% power (α = 0.05; using the “simr” package in R^116^). Specifically, we ran 500 simulations per sample size to estimate power for testing two key effects underlying the hypothesized reciprocal interplay between temporal memory and affect dynamics: 1) the association between temporal distance memory and affective spillover, and 2) the effect of emotional-valence shifts on temporal distance memory (for details, see Fig. 1). Due to the novelty of the hypothesized effects, we derived effect sizes and participant- and trial-level variance components from two most relevant studies. For 1) the association between temporal distance memory and affective spillover, these parameters came from our recent study revealing a correlation between temporal memory and emotion biases reflecting temporal neglect (see Fig. 4b in^42^; standardized *β* = 0.069, estimated from publicly available data: https://osf.io/3h8ns). For 2) the effect of emotional-valence shifts on temporal distance memory, they came from a recent study showing the effects of emotional-valence shifts on temporal memory (see Fig. 3b middle panel in^63^, Cohen’s d = 0.085; estimated from publicly available data: https://doi.org/10.17605/osf.io/s8g5n). The simulation results suggested that a sample size of N = 30 is sufficient to detect both effects (for details, see deposited code: https://osf.io/db3vq/files/osfstorage/6ae8cf6f28bb22fb03118564). Thus, our final sample size N = 51 was adequately powered to detect effect sizes reported in relevant prior studies.

Participants were recruited through flyers and the subject pool at the University of California, Santa Barbara. All participants were fluent English speakers with normal or corrected- to-normal vision and no reported history of neurological or psychiatric disorders requiring hospitalization. All procedures were approved by the Human Subjects Review Committee at the University of California, Santa Barbara, and informed consent was obtained electronically prior to participation. Participants received course credit or monetary compensation.

### Stimuli

#### Emotional images and sequences

A total of 280 emotional images (140 positive, 140 negative) were selected from well-validated databases: 157 from the International Affective Picture System (IAPS)^117^, 81 from the Open Affective Standardized Image Set (OASIS)^75^, and 42 from the Nencki Affective Picture System (NAPS)^118^. Normative valence and arousal ratings of images from the OASIS database were converted from a 1–7 scale to the 1–9 scale used in IAPS and NAPS, using the following formula: converted rating = 1 + (original rating − 1) × 8/6 (for details, see deposited code: https://osf.io/db3vq/files/osfstorage/69b1a528cfe5359581d29ec0). Positive and negative images were matched on valence extremity (absolute distance from neutrality) and arousal (valence extremity: *t* = 1.06, *p* = 0.29, *M*_Positive_ = 2.26, *SD*_Positive_ = 0.49, *M*_Negative_ = 2.20, *SD*_Negative_ = 0.56; arousal: *t* = −1.26, *p* = 0.21, *M*_Positive_ = 5.22, *SD*_Positive_ = 0.75, *M*_Negative_ = 5.33, *SD*_Negative_ = 0.73), as well as on low-level visual features, including mean luminance and visual complexity^119^ (*ps* > 0.10), quantified using a custom MATLAB (vR2021b; MathWorks) script adapted from the “lumCal” function from the SHINE color toolbox^120^. For a subset of the images, image luminance was manually adjusted using Adobe Photoshop (v2022) to better equate the average luminance across positive and negative picture sets.

Two stimulus lists were created and counterbalanced across participants, each containing 140 four-image sequences (70 positive, 70 negative). Given the limited pool of 280 images matched for emotional valence extremity and arousal, each image was used twice to create a large number (N = 140) of distinct emotional sequences to ensure adequate statistical power. Sequence construction followed several constraints: First, sequence order was pseudorandomized with no image repeated until all images had been presented once. Second, images presented in a given serial position on their first occurrence were redistributed across positions on their second occurrence. To avoid inducing systematic affective trajectories across sequences, we ensured that valence extremity and arousal did not vary systematically across the four serial positions for either positive or negative sequences, and no serial position by valence interactions were observed (*ps* > 0.10). Finally, primacy, peak, end, mean, and standard deviation of valence extremity and arousal across the four positions were matched between positive and negative sequences (*ps* > 0.10).

#### Neutral faces

A total of 140 neutral face images (half male, half female) were selected from well-established face databases, including the MacBrain Face Stimulus Set (https://macbrain.org/resources), the Karolinska Directed Emotional Faces set (https://kdef.se/home/aboutKDEF), and the Chicago Face Database^76^. All face images were cropped to minimize external features (e.g., clothing), converted to grayscale, and matched for mean luminance and root-mean-square contrast using the SHINE color toolbox^120^ in MATLAB (vR2021b) and Adobe Photoshop (v2022).

### Task design and procedures

#### Overview

Participants completed 10 runs of the experimental paradigm. Each run comprised an *Emotional Sequence Spillover Task*, in which participants viewed dynamically shifting positive and negative emotional sequences interleaved with neutral faces, followed by a *Temporal Memory Task* assessing memory for temporal distance and order of image pairs sampled from within and across emotional sequences. The two tasks were separated by a 30-s distraction task in which participants tracked the direction of a moving dot using left and right key presses (**Fig. 2a**).

#### Emotional sequence spillover task

In each run, participants viewed 14 emotional sequences (7 positive, 7 negative), each comprising four emotional images. Following each sequence, a neutral face was presented (50% female; balanced across valences), and participants were asked to rate it based on likeability using a continuous scale ranging from 1 (“*do not like/trust at all*”) to 4 (“*like/trust very much*”) (**Fig. 2b**). Face-likeability ratings indexed affective spillover—i.e., the extent to which the preceding emotional sequences biased face likeability in a valence-congruent manner (as indexed by higher likeability following positive sequences and lower likeability following negative sequences). Following previous work^9^, to minimize demand characteristics, participants were told that the study examined how people form first impressions of others in the presence of emotional distractors. They were instructed to base their likeability ratings on their immediate, intuitive impressions of each face, focusing on perceived trustworthiness (e.g., whether they would feel comfortable asking the person for help) rather than attractiveness. Emotional images were described as task-irrelevant distractors to prevent participants from intentionally basing their face ratings on the immediately preceding emotional content. Participants were fully debriefed upon completion of the study.

To facilitate temporal memory encoding, participants were encouraged to form a brief story linking each pair of adjacent emotional images^121,122^. Each image was presented for 2 s with a visual angle of 9.6° × 7.2°, followed by a fixation screen jittered between 1.6 and 2 s. Each face was presented for 0.45 s at a visual angle of 5.7° × 7.2°, followed by a 2-s rating screen and then a jittered fixation interval (1.15–1.55 s) before the onset of the next sequence.

#### Emotional-valence transitions

We systematically manipulated transitions between emotional sequences to produce shifts both within and across valence. Sequence presentation order was pseudorandomized for each participant such that, within each run, the N = 13 transitions between N = 14 sequences included approximately equal numbers of each transition type. Specifically, across the 10 experimental runs, 13 transitions in half the runs included 3 positive-to-positive, 3 negative-to-negative, 3 positive-to-negative, and 4 negative-to-positive, while the other half included 3 positive-to-positive, 3 negative-to-negative, 4 positive-to-negative, and 3 negative-to-positive. We further ensured that no more than two sequences of the same valence occurred consecutively. Such participant-specific pseudorandomization also ensured that valence transitions were decoupled from transitions between specific items—for instance, a positive-to-negative transition could be generated by a birthday cake-barking dog pair in one participant but a birthday cake-car accident pair in another. This reduces the possibility that the observed valence-transition effects could be attributed to the idiosyncratic properties of particular item pairs.

#### Temporal memory task

After every 14 emotional sequences (and an intervening distraction task), participants completed a temporal memory task assessing temporal order and distance memory for image pairs drawn from the just-viewed sequences, including N = 14 within-sequence pairs and N = 13 across-sequence pairs (corresponding to the N = 13 unique transitions between the N = 14 sequences). On each trial, two images were displayed simultaneously to the left and right of a central fixation cross. Participants were asked to indicate which image they thought had occurred earlier (i.e., indexing *temporal order memory*). Next, they were asked to indicate the remembered temporal distance between the two images on a continuous scale ranging from “*very close*” to “*very far*” (i.e., indexing *temporal distance memory*) (**Fig. 2c**). Responses were self-paced, and participants were encouraged to respond as accurately and quickly as possible (order memory: *M*_RT_ = 5.47, *SD*_RT_ = 1.42; distance memory: *M*_RT_ = 1.36; *SD*_RT_ = 0.48, in seconds). Left–right image placement on the screen was counterbalanced such that the earlier-occurring image was equally likely to appear on either side.

#### Within-vs. across-sequence pairs

Unbeknownst to the participants, within-sequence pairs consisted of the second and the last images of the same sequence, whereas across-sequence pairs consisted of the last image of one sequence and the first image of its following sequence. Because each emotional image occurred twice during encoding, and the last image of each sequence was included in both within-and across-sequence pairs, individual images could be probed multiple times during temporal memory testing (for control analyses accounting for item exposure history in our primary temporal memory analyses, see *Temporal memory: Item exposure history in Supplementary Information*). Within- and across-sequence pairs were matched on objective temporal distance (i.e., time elapsed between the onset of the two images during encoding; *M*_Within_ = 7.6s, *M*_Across_ = 7.6s; *p*s > 0.10), as well as on emotional properties (valence extremity and arousal) and low-level visual features (luminance and complexity) (*p*s > 0.10).

### EEG recording and preprocessing

EEG was continuously recorded throughout the task using a 64-channel Brain Products actiCHamp system, sampled at 500Hz and referenced online to Cz. Electrode impedances were kept below 10 kΩ. Eye movements and blinks were monitored using electrooculogram (EOG) electrodes placed at the lateral canthi (horizontal EOG) and above and below the right eye (vertical EOG). Channels exhibiting excessive noise, likely due to poor scalp contact, were visually identified and interpolated (on average, 0.88 ± 1.10 of 64 channels per run per participant). Independent Component Analysis (ICA) was performed for artefact detection. Components were visually inspected, and those reflecting eye blinks and eye movements were removed (4.71 ± 2.69 of 65 components per participant). In limited cases, components reflecting channel noise— characterized by a highly focal scalp topography centered on a single electrode and confirmed by visual inspection of the corresponding channel time series—were removed (0.31 ± 0.71 of 65 components per participant). Data were then re-referenced to the average of all scalp electrodes.

Preprocessing steps were performed using EEGLAB (v2022) functions123 and custom MATLAB (vR2021b; MathWorks) scripts.

### Alpha burst detection

We estimated nonstationary alpha-band (8–12 Hz) oscillatory bursts to examine their contribution to online temporal memory coding during emotional sequences. Continuous EEG data from each sequence onset to the end of the post-sequence interval were subjected to the cycle-by-cycle analysis^73^ (amplitude fraction threshold = 0.3; amplitude consistency threshold = 0.4; period consistency threshold = 0.4; monotonicity threshold = 0.8; minimum cycles = 3)^66,67,78^. For each electrode, we computed *alpha burst time* as the proportion of time during which alpha activities were classified as oscillatory bursts, which was estimated separately for image-presentation and ISI epochs. Stimulus-induced alpha burst time was then averaged across a predefined cluster of parieto-occipital electrodes (POz, PO3, PO4, PO7, PO8, O1, O2, & Oz^66,67,77–79^) and baseline-corrected by subtracting trial-averaged ISI burst time. For across-sequence analyses, alpha burst time was averaged across the two images in each pair and baseline-corrected using trial-averaged burst time during the post-image intervals. Cycle-by-cycle analysis was not applied to the intervening face epochs because their short duration (0.45 s) precluded reliable burst detection. Finally, to assess the potential process specificity of alpha bursts to temporal memory indices, identical procedures (with different parameters) were applied to the beta frequency band for control analyses (see *Cycle-by-cycle analysis: Beta bursts* in *Supplementary Methods*).

### Alpha center power estimation

To examine whether burst-related effects are specific to nonstationary alpha activity rather than reflecting the effects of sustained oscillatory alpha power, we additionally extracted trial-wise alpha power at the center frequency (i.e., alpha center power) as follows. Continuous EEG data were epoched from −1000 to 2500 ms relative to emotional-image onset. Epochs containing remaining artifacts, identified via visual inspection, were excluded (on average, 0.63 ± 0.84% per subject). Time–frequency decomposition was performed on single-trial epochs using complex Morlet wavelets (1–50 Hz in 0.5-Hz steps; 3–10 cycles)^78^ using a custom MATLAB script adapted from [publicly available code](https://drive.google.com/a/ucsc.edu/file/d/174f2JT_T9kSeXSGl2qcqqRIltIQ8brnR/view?usp=sharing). Power spectra for emotional images were computed by averaging power across time points within a pre-image baseline window (−500 to −300 ms) and the image-presentation window (0–2000 ms). Power spectra from each window, channel, and participant were parameterized using the fitting oscillations and one-over-f (FOOOF) algorithm^80^ over 1–50 Hz (peak width limits = 1–8 Hz; maximum number of peaks = 6; minimum peak height = 0.1; peak threshold = 1.5; aperiodic mode = fixed^78^). Alpha center power was averaged across a predefined cluster of parieto-occipital electrodes (POz, PO3, PO4, PO7, PO8, O1, O2, & Oz^66,67,77–79^) and baseline-corrected by subtracting the across-trial average baseline. Alpha center frequency was additionally extracted for control analyses. For further details, see *Validation of alpha center power estimation* in *Supplementary Results*.

### Statistical analysis

*Mixed-effects modeling*. Analyses were conducted using linear mixed-effects models (for continuous outcome variables) and generalized linear mixed-effects models (for binary outcome variables) using the “lme4” package in R^124^, unless otherwise specified. Predictor variables were included as fixed effects. All models included by-participant random intercepts. Models predicting face-likeability ratings and affective spillover additionally included by-face random intercepts to account for face-specific variance. Models predicting within-sequence temporal memory additionally included by-sequence random intercepts. By-participant, by-face, or by-sequence random slopes were further included when they: (a) did not produce convergence errors, and (b) significantly improved model fit relative to random-intercept-only models, as assessed by likelihood-ratio tests^125^ and the Bayesian Information Criterion (BIC). When convergence issues occurred, we attempted to resolve them using an alternative optimizer (“bobyqa”) in the “lmer” function^126^; if convergence could not be achieved, the corresponding random slopes were excluded. Full model descriptions for each primary model, including the final random-effects structure, are reported in *Supplementary Information: Mixed-effects modeling: Random-effects structure for primary analyses*. Multiple-comparison corrections were applied using the false discovery rate (FDR) approach^127^ (for details, see *Neural correlates of temporal memory coding and affect dynamics: Alpha-burst analysis*). The stimulus list (see *Emotional images and sequences* above) was included as a fixed-effect control factor in all mixed-effects models.

### Primary analyses

#### Affective spillover

To test for affective spillover, we examined whether emotional-sequence valence (categorical: positive *vs.* negative; used throughout) predicted likeability ratings of subsequently presented neutral faces. We further tested whether this effect remained significant after controlling for normative face-likeability ratings and face gender (see *Affective spillover: Face characteristics* in *Supplementary Results*). Normative face-likeability ratings were obtained from an independent sample (N = 100) who rated the neutral faces in the absence of emotional sequences. We also computed a *trialwise* affective spillover metric, which was used for examining the associations between spillover and temporal distance and order memory (see below). This metric was calculated as the difference between a participant’s in-task rating and the normative baseline rating of each face, signed according to the valence of the preceding sequence, such that positive values reflected affective spillover (i.e., ratings higher than baseline following positive sequences and lower than baseline following negative sequences).

#### Emotional-sequence boundary effects on temporal memory

We first tested whether temporal order memory accuracy was significantly above chance across participants, using a one-sample *t* test against chance (50%). To examine event-boundary effects induced by transitions of emotional sequences (regardless of emotional-valence shifts), we tested whether trialwise temporal distance ratings and order memory accuracy were modulated by image-pair type (sampled within *vs.* across sequences). To ensure that any across-*vs.* within-sequence differences were not solely driven by emotional-valence shifts across sequences, we repeated the analyses restricting across-sequence pairs to those belonging to the same valence; we also tested whether emotional-sequence valence interacted with image-pair type to predict temporal distance ratings and order memory.

#### Within-sequence temporal memory and affective spillover

To examine whether emotional events associated with fine-grained, high-fidelity temporal memory were associated with subsequent affective spillover, we tested whether trialwise temporal distance ratings and order memory for within-sequence image pairs predicted spillover produced by those sequences. We also tested whether emotional-sequence valence moderated these associations.

#### Emotional-valence shifts and across-sequence temporal memory

To examine the influence of emotional-valence shifts on temporal memory, we tested whether temporal distance and order memory across sequences were modulated by the interaction between the valence of each sequence and that of the immediately following sequence (first *vs.* second valence interaction). When this interaction was significant, we conducted a planned contrast analysis comparing image pairs that spanned a valence shift (positive-to-negative or negative-to-positive) with those that did not (positive-to-positive or negative-to-negative). To assess potential directional effects of emotional-valence shifts, we further contrasted positive-to-negative and negative-to-positive pairs.

When a significant directional difference in order memory was observed, we conducted signal detection analyses^128^ to determine whether it reflected differences in sensitivity or response bias. Trials in which negative images truly occurred later (positive-to-negative pairs) were treated as “signal.” For each participant, we calculated sensitivity (*d*′ = z(Hit) − z(False Alarm)), indexing discrimination of the correct temporal order, and response bias (C = −0.5 × [z(Hit) + z(False Alarm)]), indexing the tendency to judge negative images as occurring later regardless of accuracy. We then tested whether *d*′ and C differed significantly from zero across participants, as evidence of sensitivity or bias toward remembering negative images as later.

#### Neural correlates of temporal memory coding and affect dynamics: Alpha-burst analysis

To examine the contribution of alpha bursts to online temporal memory coding, we tested whether baseline-corrected, trialwise alpha burst time during emotional image viewing predicted temporal distance and temporal order memory for image pairs within the corresponding sequences. To test whether associations with temporal memory indices were specific to stimulus-evoked alpha bursting activity during image encoding, we conducted parallel analyses using alpha burst time extracted from the ISIs within emotional sequences. To determine whether burst-related effects were specific to nonstationary alpha activity rather than reflecting the effects of sustained oscillatory alpha power, we additionally tested whether alpha burst time predicted temporal distance and order memory while controlling for alpha center power. Finally, to establish feature and frequency specificity, we examined whether alpha center frequency as well as burst time in the beta band, predicted temporal distance and temporal order memory within emotional sequences (see *Within-sequence temporal memory: Specificity of alpha-burst effects* in *Supplementary Results*). FDR correction was applied across analyses examining multiple alpha measures (burst time, center power, and center frequency), frequency bands (burst time in the alpha and beta bands), as well as timing windows (alpha burst time in image encoding vs. ISIs). To examine whether alpha burst time was associated with affective spillover, we tested whether baseline-corrected, trialwise alpha burst time during emotional image viewing predicted subsequent affective spillover, and whether these associations differed by emotional-sequence valence. We additionally tested whether alpha burst time itself varied as a function of emotional-sequence properties (i.e., valence and normative arousal). Parallel analyses were conducted for alpha center power to evaluate whether any observed associations with affective spillover were specific to alpha burst dynamics rather than sustained oscillatory power, and FDR correction was applied across analyses of alpha burst time and center power (see *Within-sequence temporal memory and affective spillover: Alpha center power* in *Supplementary Results*). Of note, trials with alpha burst time or alpha center power exceeding 4 SDs from the mean across trials within each subject were excluded from the relevant analyses (1.60 ± 1.37% and 1.27 ± 1.44% of trials, respectively).

To assess whether alpha bursts also contributed to temporal memory coding across sequences, we tested whether alpha burst time elicited during the encoding of images from across-sequence pairs (see *Alpha burst detection* in *Methods*) predicted temporal distance and temporal order memory. We additionally examined whether alpha burst time varied as a function of emotional-valence transitions across sequences. Detailed results are reported in *Across-sequence temporal memory: Alpha modulation* in *Supplementary Information*.

### Control analyses

We conducted additional control analyses to assess whether the associations between temporal memory and affective spillover, as well as the effects of emotional-valence transitions on temporal memory, remained robust after accounting for additional factors, including arousal^11,14,19,36,110^, semantic similarity^32,81–84^, and item exposure history during temporal memory testing. We also examined the independent effects of these factors on temporal memory. Detailed results are reported in *Supplementary Information*.

## Data and code availability

All data reported here, as well as the scripts used to run the experiment and perform the analyses, are available at: https://osf.io/db3vq.

## Acknowledgments

This work is supported by the National Institute of Mental Health grant R01-MH134000 (R. C. L.) and by the Academic Senate at the University of California, Santa Barbara (R. C. L.). The authors thank L. Azizi, J. Samaha, T. Sprague, and J. Wang for helpful discussions, and R. Wang, J. Diaz, J. Sun, J. Adda, H. Lu, and A. Ayesh for assistance with data collection and/or EEG data preprocessing. Author contributions: conceptualization and methodology, M.L. and R.C.L.; investigation, M.L., B.S., and T.N.L.; project administration, M.L., B.S., and T.N.L.; formal analysis, M.L., R.C.L., and B.G.; visualization, M.L. and R.C.L.; writing – original draft, M.L. and R.C.L.; writing – review and editing, M.L., R.C.L., B.S., T.N.L., and B.G.; data curation, M.L.; resources, R.C.L.; supervision, B.G. and R.C.L.; and funding acquisition, R.C.L.

