## Supplementary Information for "The temporal organization of memory and emotion is reciprocally coupled"

**Acknowledgments**

This work is supported by the National Institute of Mental Health grant R01-MH134000 (R. C. L.) and by the Academic Senate at the University of California, Santa Barbara (R. C. L.). The authors thank L. Azizi, J. Samaha, T. Sprague, and J. Wang for helpful discussions, and R. Wang, J. Diaz, J. Sun, J. Adda, H. Lu, and A. Ayesh for assistance with data collection and/or EEG data preprocessing. Author contributions: conceptualization and methodology, M.L. and R.C.L.; investigation, M.L., B.S., and T.N.L.; project administration, M.L., B.S., and T.N.L.; formal analysis, M.L., R.C.L., and B.G.; visualization, M.L. and R.C.L.; writing – original draft, M.L. and R.C.L.; writing – review and editing, M.L., R.C.L., B.S., T.N.L., and B.G.; data curation, M.L.; resources, R.C.L.; supervision, B.G. and R.C.L.; and funding acquisition, R.C.L.

**Conflict of Interest Statement:** The authors declare no competing financial interests.

### Supplementary Methods

*Cycle-by-cycle analysis: Beta bursts.* The following parameters were used for cycle-by-cycle analysis in the beta frequency band (13-30 Hz): amplitude fraction threshold = 0.4; amplitude consistency threshold = 0.5; period consistency threshold = 0.5; monotonicity threshold = 0.85; minimum cycles = 3. These criteria were more stringent than those applied to the alpha band because beta oscillations are faster and shorter in duration, requiring stricter thresholds to avoid classifying transient or noise-driven fluctuations as bursts[1,2].

### Supplementary Results

*Affective spillover: Face characteristics.* Beyond emotional-sequence valence, both normative likeability ($B = 1.27$ ($SE = 0.05$), 95% CI [1.18, 1.37], $F = 714.01$, $p < 0.001$) and neutral-face gender significantly predicted face likeability ratings in the task ($F = 21.55$, $p < 0.001$). Female faces (*vs.* male faces) and faces with higher normative likeability ratings received higher likeability ratings in the task ($B = 0.11$ ($SE = 0.02$), 95% CI [0.07, 0.16], $t = 4.64$, $p < 0.001$, $M_{\text{Female}} = 2.64$, $SE_{\text{Female}} = 0.03$, $M_{\text{Male}} = 2.26$, $SE_{\text{Male}} = 0.03$).

*Validation of alpha center power estimation.* To confirm the presence of alpha oscillations in our data, we first inspected the time–frequency spectrum averaged across all trials and participants over a predefined parieto-occipital cluster, spanning both the pre-stimulus baseline and image-presentation periods (**Fig. S1a**). Visual inspection revealed a clear modulation of power within the alpha band, characterized by a reduction in power shortly after stimulus onset, consistent with canonical alpha event-related desynchronization. We next computed group-averaged power spectra for emotional-image trials by averaging power across time points within the pre-image baseline window (−500 to −300 ms) and the image-presentation window (0–2000 ms). These spectra were parameterized using the FOOOF algorithm[3] over 1–50 Hz. As shown in **Fig. S1b–c**, the models identified robust alpha peaks at the group level during both the baseline and stimulus windows (baseline: CF = 11.30 Hz, $R^2 = 0.987$; stimulus: CF = 10.95 Hz, $R^2 = 0.995$), with a reduced alpha peak during image presentation, consistent with stimulus-induced alpha desynchronization. Weak beta-band peaks were also identified (baseline: CF = 19.35 Hz; stimulus: CF = 19.14 Hz).

Importantly, for the primary analyses of alpha center power (see *Methods: Alpha center power estimation* in the main text), FOOOF parameterization was performed at the *single-trial* level. Across trials, alpha peaks were detected in 98.15 ± 4.28% of trials (averaged across participants), with robust model fits ($R^2 = 0.95 \pm 0.04$). Beta peaks were also detected in 99.52 ± 1.19% of trials.

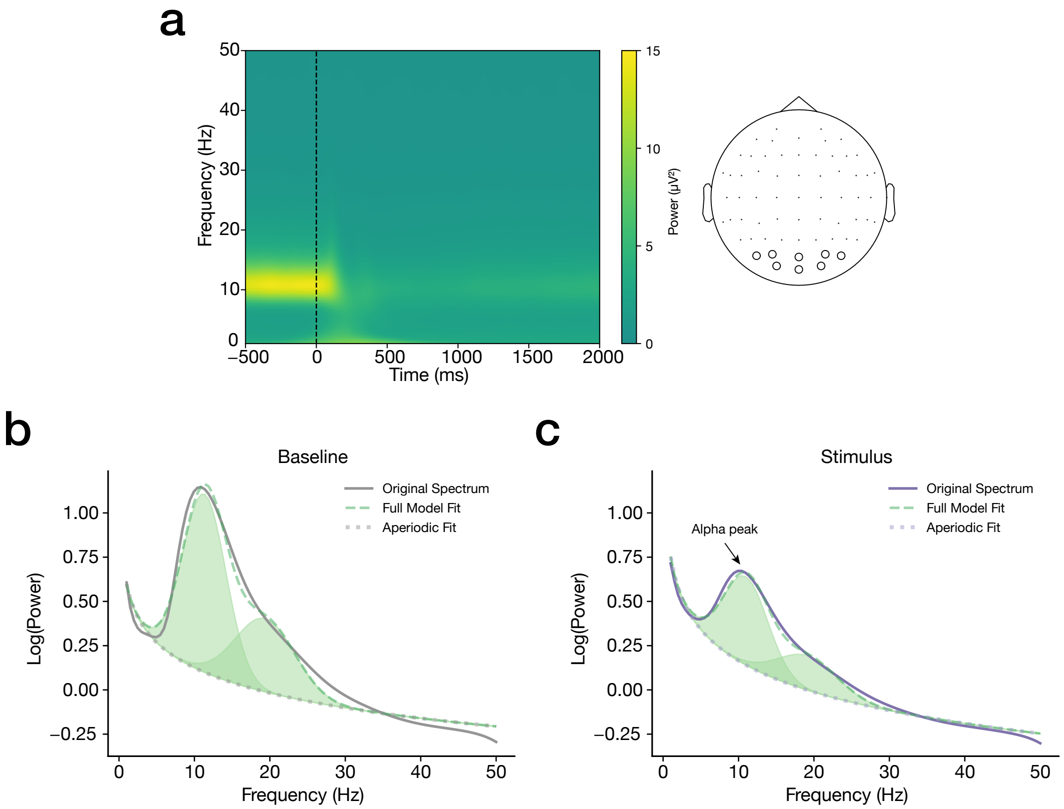


**Fig. S1. Validation of alpha center power estimation. a)** Time–frequency representation (1–50 Hz) averaged over the parieto-occipital cluster across all trials and subjects, time-locked to emotional-image onset (0 ms; dashed line). Warmer colors indicate greater power. Visual inspection revealed a clear power modulation within the alpha band (8–12 Hz), characterized by a reduction in power shortly after stimulus onset, consistent with canonical alpha event-related desynchronization. Group-averaged power spectra (solid line) during **b)** the pre-stimulus baseline (−500 to −300 ms) and **c)** image-presentation windows (0–2000 ms) were parameterized with FOOOF, decomposed into the aperiodic (dotted line) and periodic components; shaded regions indicate detected oscillatory peaks. These parameterizations demonstrate reliable alpha peaks during both the baseline and stimulus windows (baseline: CF = 11.30 Hz, $R^2 = 0.987$; stimulus: CF = 10.95 Hz, $R^2 = 0.995$), with a reduced alpha peak during image presentation. Weaker peaks were also detected in the beta band. Of note, for the primary analyses of alpha center power (see *Methods: Alpha center power estimation* in the main text), FOOOF parameterization was performed at the single-trial level.

*Within-sequence temporal memory and affective spillover: Alpha center power.* We examined whether alpha center power during emotional-sequence encoding correlated with temporal distance and order memory, and whether the effects of alpha burst time remained significant after controlling for center power, suggesting the specific contribution of nonstationary rather than sustained oscillatory alpha activity to temporal memory coding. Similar to greater alpha burst time, greater alpha center power during emotional sequences predicted longer remembered temporal distance for image pairs within the sequences (**Fig. S2a**; $B = 0.89$ ($SE = 0.15$), 95% CI [0.59, 1.19], $F = 33.74$, $p_{FDR} < 0.001$), as well as poorer order memory (**Fig. S2b**; $B = -1.46$ ($SE = 0.23$), 95% CI [−1.90, −1.02], OR = 0.23, 95% CI [0.15, 0.36], $\chi^2(1) = 41.78$, $p_{FDR} < 0.001$). Critically, alpha burst time continued to significantly predict remembered temporal distance even after controlling for alpha center power (burst time: $B = 1.80$ ($SE = 0.52$), 95% CI [0.78, 2.82], $F = 11.98$, $p = 0.0006$; center power: $B = 0.18$ ($SE = 0.23$), 95% CI [−0.28, 0.64], $F = 0.59$, $p = 0.44$). Interestingly, however, temporal order memory was instead most reliably associated with reduced alpha center power rather than alpha burst time, in a simultaneous model (center power: $B = -1.16$ ($SE = 0.34$), 95% CI [−1.83, −0.48], OR = 0.31, 95% CI [0.16, 0.62], $\chi^2(1) = 11.31$, $p = 0.0008$; burst time: $B = -0.45$ ($SE = 0.72$), 95% CI [−1.86, 0.96], OR = 0.64, 95% CI [0.16, 2.61], $\chi^2(1) = 0.39$, $p = 0.53$). Together, these findings suggest that alpha burst dynamics and alpha power reflect partially dissociable mechanisms supporting the temporal organization of emotional experiences in memory, with alpha bursts more closely tracking temporal distance coding.

We additionally examined whether alpha center power was associated with affective spillover, and tested whether such an association differed by sequence valence. Similar to alpha burst time, alpha center power significantly interacted with valence in predicting affective spillover (**Fig. S2c**; $F = 62.92$, $p_{FDR} < 0.001$), such that greater alpha power during negative sequences was associated with reduced subsequent spillover ($B = -0.38$ ($SE = 0.08$), 95% CI [−0.53, −0.22], $t = -4.78$, $p < 0.001$), whereas greater alpha power during positive sequences was associated with increased spillover ($B = 0.35$ ($SE = 0.08$), 95% CI [0.20, 0.51], $t = 4.40$, $p < 0.001$). We further tested whether alpha center power itself was sensitive to sequence valence or arousal, and found that higher normative arousal was associated with lower alpha power (i.e., greater event-related desynchronization; $B = -0.009$ ($SE = 0.004$), 95% CI [−0.02, −0.002], $F = 5.91$, $p_{\mathrm{FDR}} = 0.031$; **Fig. S2d**), consistent with prior findings[4–7]. Together, these findings suggest that alpha center power may index arousal-related encoding processes that contribute to temporal memory organization and modulate affective persistence in a valence-dependent manner.

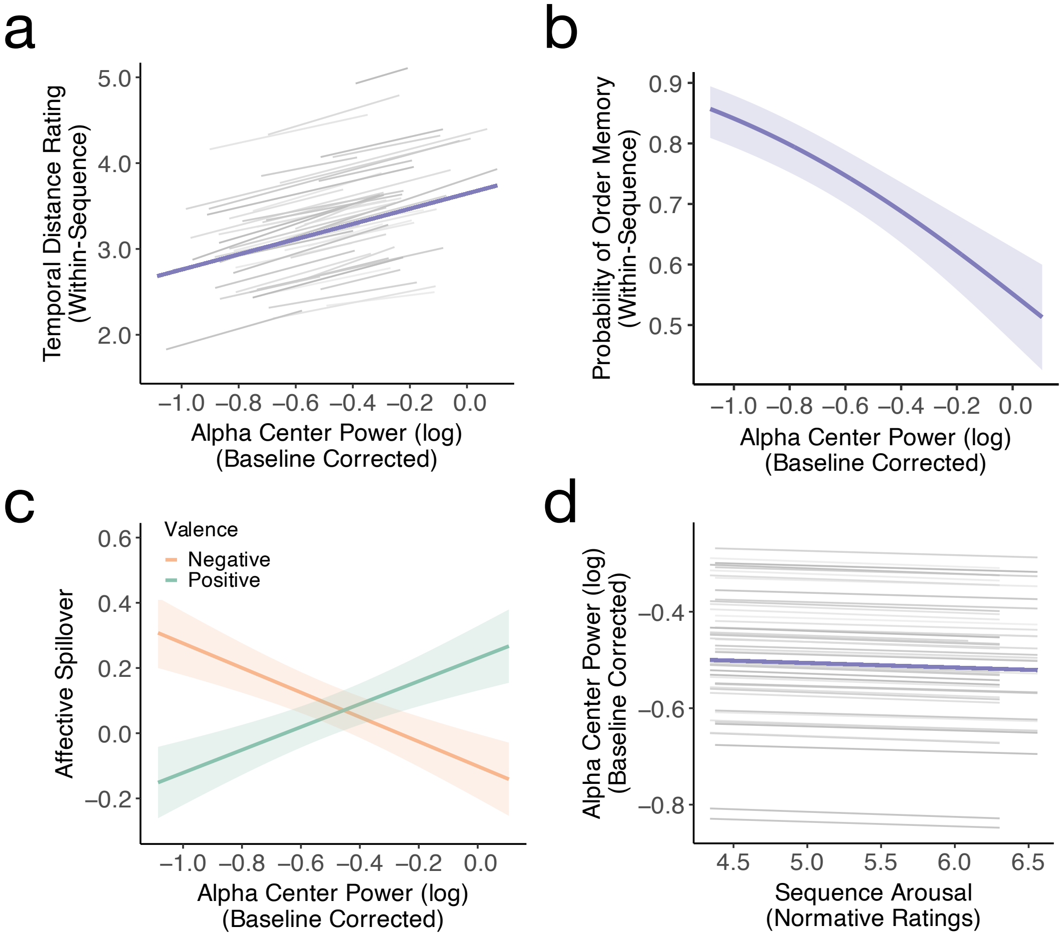


**Fig. S2. Alpha center power contributes to online temporal memory coding and subsequent affective spillover**. **a)** Greater alpha center power during emotional sequences predicted longer remembered temporal distance for image pairs within the sequences ($B = 0.89$, $p_{FDR} < 0.001$), as well as **b)** poorer order memory ($\chi^2(1) = 41.78$, $p_{\mathrm{FDR}} < 0.001$). **c)** Alpha center power significantly interacted with valence in predicting affective spillover ($F = 62.92$, $p_{\mathrm{FDR}} < 0.001$), such that greater alpha power during negative sequences was associated with reduced subsequent spillover ($B = -0.38$, $p < 0.001$), whereas greater alpha power during positive sequences was associated with increased spillover ($B = 0.35$, $p < 0.001$). Finally, sequences with higher normative arousal induced lower alpha power ($B = -0.009$, $p_{\mathrm{FDR}} = 0.031$).

*Within-sequence temporal memory: Specificity of alpha-burst effects.* To further assess the specificity of the observed alpha-burst effects on temporal memory, we examined whether alpha center frequency and burst time in the beta band were associated with within-sequence temporal distance and order memory. None of these measures significantly predicted either temporal distance or temporal order memory (alpha center frequency predicting distance memory: $B = 0.05$ ($SE = 0.04$), 95% CI [−0.03, 0.13], $F = 1.66$, $p_{FDR} = 0.20$, and order memory: $B = 0.09$ ($SE = 0.06$), 95% CI [−0.02, 0.21], OR = 1.10, 95% CI [0.98, 1.24], $\chi^2(1) = 2.41$, $p_{FDR} = 0.14$; beta burst time predicting distance memory: $B = 2.19$ ($SE = 1.20$), 95% CI [−0.17, 4.54], $F = 3.12$, $p_{FDR} = 0.10$, and order memory: $B = -1.78$ ($SE = 1.57$), 95% CI [−4.87, 1.31], OR = 0.17, 95% CI [0.008, 3.69], $\chi^2(1) = 1.28$, $p_{FDR} = 0.26$). Together, these results support the feature- and frequency-specific nature of the reported alpha-burst effects.

*Within-sequence temporal memory and affective spillover: Arousal effects*. We tested whether the association between within-sequence temporal (distance and order) memory and affective spillover was moderated by sequence-level normative arousal (averaged across images) or by the valence × arousal interaction. Arousal did not significantly moderate these associations (distance memory: $F = 0.31$, $p = 0.58$; order memory: $F = 0.01$, $p = 0.91$). The three-way interaction between temporal (distance or order) memory, valence, and arousal was also not significant (distance memory: $F = 1.18$, $p = 0.28$; order memory: $F = 0.27$, $p = 0.61$).

*Across-sequence temporal memory: Arousal-change effects.* The effects of emotional-valence transitions on temporal distance and order memory across sequences remained significant after controlling for signed arousal differences between the two images in each across-sequence pair (image 2 *vs.* 1 arousal; distance memory: $F = 184.16$, $p < 0.001$; order memory: $\chi^2(1) = 19.08$, $p < 0.001$). In contrast, arousal differences themselves did not significantly predict either measure of temporal memory (distance memory: $B = -0.005$ ($SE = 0.02$), 95% CI [−0.04, 0.03], $F = 0.08$, $p = 0.77$; order memory: $B = 0.04$ ($SE = 0.02$), 95% CI [−0.001, 0.09], OR = 1.05, 95% CI [1.00, 1.09], $\chi^2(1) = 3.60$, $p = 0.06$).

We further tested whether graded emotional changes across sequences predicted temporal memory by modeling the signed change in normative valence and arousal between the two images in each pair, as well as their interaction. Analyses were conducted separately for same-valence and different-valence pairs. For pairs that differed in valence, *shifting toward a more negative image* (i.e., more negative Δ valence) predicted better temporal order memory ($B = 0.02$, $p = 0.02$), consistent with the categorical valence-transition effects reported in the main text (i.e., better order memory for *positive-to-negative* relative to negative-to-positive transitions). No other effects reached significance ($p$s > 0.05). See Supplementary Table 1 below for full model statistics.

**Supplementary Table 1**
*Mixed-effects modeling results on the effects of signed valence and arousal differences on across-sequence temporal memory*

| Outcome and predictor | $B$ ($SE$) | 95% CI | OR [95% CI] | Test | $p$ |
|---|---|---|---|---|---|
| **Same-valence pairs** | | | | | |
| ***Temporal distance memory*** | | | | | |
| Δ Valence | −0.01 (0.03) | [−0.07, 0.06] | — | $F = 0.03$ | 0.87 |
| Δ Arousal | −0.02 (0.02) | [−0.06, 0.03] | — | $F = 0.53$ | 0.47 |
| Δ Valence × Δ Arousal | −0.06 (0.03) | [−0.12, 0.001] | — | $F = 3.68$ | 0.06 |
| ***Temporal order memory*** | | | | | |
| Δ Valence | −0.04 (0.05) | [−0.14, 0.06] | 0.96 [0.87, 1.06] | $\chi^2(1) = 0.72$ | 0.40 |
| Δ Arousal | 0.06 (0.03) | [−0.01, 0.13] | 1.06 [0.99, 1.14] | $\chi^2(1) = 3.13$ | 0.08 |
| Δ Valence × Δ Arousal | −0.01 (0.04) | [−0.09, 0.08] | 0.99 [0.91, 1.09] | $\chi^2(1) = 0.01$ | 0.91 |
| **Different-valence pairs** | | | | | |
| ***Temporal distance memory*** | | | | | |
| Δ Valence | 0.004 (0.005) | [−0.005, 0.01] | — | $F = 0.81$ | 0.37 |
| Δ Arousal | −0.002 (0.02) | [−0.04, 0.04] | — | $F = 0.01$ | 0.91 |
| Δ Valence × Δ Arousal | 0.006 (0.005) | [−0.003, 0.02] | — | $F = 1.95$ | 0.16 |
| ***Temporal order memory*** | | | | | |
| Δ Valence | −0.02 (0.008) | [−0.03, −0.003] | 0.98 [0.97, 1.00] | $\chi^2(1) = 5.62$ | 0.02* |
| Δ Arousal | 0.04 (0.03) | [−0.03, 0.10] | 1.04 [0.97, 1.10] | $\chi^2(1) = 1.19$ | 0.27 |
| Δ Valence × Δ Arousal | −0.005 (0.007) | [−0.02, 0.009] | 0.99 [0.98, 1.01] | $\chi^2(1) = 0.51$ | 0.47 |

*Note.* Δ = signed difference between the two items of a pair (image 2 − image 1). Temporal distance memory was modeled with linear mixed-effects models ($F$ tests); temporal order memory was modeled with logistic mixed-effects models (Wald $\chi^2$ tests). OR = odds ratio. 95% CIs are Wald intervals. *$p < 0.05$.

*Across-sequence temporal memory: Alpha modulation.* We tested whether alpha burst time elicited during encoding of images from across-sequence pairs (averaged across the two images in each pair) predicted temporal distance or temporal order memory. No significant associations were observed for alpha burst time (distance memory: $B = 0.33$ ($SE = 0.27$), 95% CI [−0.21, 0.87], $F = 1.46$, $p = 0.23$; order memory: $B = -0.33$ ($SE = 0.34$), 95% CI [−1.00, 0.34], OR = 0.72, 95% CI [0.37, 1.41], $\chi^2(1) = 0.93$, $p = 0.34$) or alpha center power (distance memory: $B = 0.18$ ($SE = 0.13$), 95% CI [−0.08, 0.44], $F = 1.92$, $p = 0.17$; order memory: $B = -0.11$ ($SE = 0.18$), 95% CI [−0.46, 0.25], OR = 0.90, 95% CI [0.63, 1.28], $\chi^2(1) = 0.34$, $p = 0.56$). We further examined whether across-sequence alpha burst time or center power varied as a function of emotional-valence transitions (first-sequence valence × second-sequence valence interaction). Again, no significant effects were obtained for either alpha burst time ($F = 0.03$, $p = 0.87$) or alpha center power ($F = 0.03$, $p = 0.85$). These null findings may originate from the complex structure at sequence boundaries—namely, transitions between emotional sequences accompanied by the presentation and evaluation of intervening neutral faces—which likely engaged a mixture of cognitive and affective processes that obscured or counteracted the contribution of alpha dynamics to temporal memory coding across sequences.

*Temporal memory: Emotional modulation*. Given growing evidence that emotional valence and/or arousal can influence temporal memory (for reviews, see[8–10]), we examined whether sequence valence, arousal, or their interaction predicted temporal distance and order memory *within* sequences, and whether average valence, arousal, or their interaction of image pairs *across* sequences predicted temporal distance and order memory. We only found that across-sequence image pairs with higher arousal were associated with poorer temporal order memory ($B = -0.13$, $p = 0.008$), consistent with prior work[11,12] (but see[13,14]). All other associations were non-significant ($p$s > 0.06). Importantly, the associations between temporal (distance and order) memory and affective spillover, as well as the effects of emotional-valence transitions on temporal memory (distance and order), remained significant after controlling for valence, arousal, or their interaction ($p$s < 0.05). See Supplementary Tables 2 and 3 below for full model statistics.

*Temporal memory: Semantic similarity.* In light of evidence that semantic relatedness between items can shape both item memory[15–18] and temporal memory[19], we examined whether semantic similarity shaped temporal memory within or across sequences. Semantic similarity was estimated using the deep convolutional neural network VGG-16[20]. Activation patterns from this network's "fc2" layer have been shown to align with representations in inferior temporal cortex that support object categorization and semantic judgements (e.g.[21–24]); thus, similarity between its activation patterns serves as an index of semantic similarity. Following recent work[25], we extracted VGG-16 activation vectors for each image, applied principal component analysis to reduce dimensionality, and computed pairwise Pearson's correlations between activation vectors for all image pairs.

**Supplementary Table 2**
*Mixed-effects modeling results on the effects of valence and arousal on temporal memory*

| **Outcome and predictor** | ***B* (*SE*)** | **95% CI** | **OR [95% CI]** | **Test** | ***p*** |
|---|---|---|---|---|---|
| **Within-sequence temporal memory** | | | | | |
| ***Temporal distance memory*** | | | | | |
| Sequence valence (positive vs. negative) | 0.01 (0.04) | [−0.07, 0.10] | — | $F = 0.09$ | 0.76 |
| Sequence arousal | −0.02 (0.06) | [−0.13, 0.09] | — | $F = 0.12$ | 0.73 |
| Sequence valence × Sequence arousal | 0.10 (0.11) | [−0.12, 0.32] | — | $F = 0.81$ | 0.37 |
| ***Temporal order memory*** | | | | | |
| Sequence valence (positive vs. negative) | −0.09 (0.06) | [−0.22, 0.03] | 0.91 [0.81, 1.03] | $\chi^2(1) = 2.36$ | 0.12 |
| Sequence arousal | 0.08 (0.08) | [−0.08, 0.23] | 1.08 [0.93, 1.26] | $\chi^2(1) = 0.94$ | 0.33 |
| Sequence valence × Sequence arousal | 0.02 (0.16) | [−0.29, 0.32] | 1.02 [0.75, 1.38] | $\chi^2(1) = 0.01$ | 0.92 |
| **Across-sequence temporal memory** | | | | | |
| ***Temporal distance memory*** | | | | | |
| Average pair valence | −0.0003 (0.01) | [−0.02, 0.02] | — | $F = 0.00$ | 0.98 |
| Average pair arousal | −0.02 (0.04) | [−0.09, 0.05] | — | $F = 0.31$ | 0.58 |
| Average pair valence × Average pair arousal | 0.04 (0.02) | [−0.002, 0.08] | — | $F = 3.44$ | 0.06 |
| ***Temporal order memory*** | | | | | |
| Average pair valence | 0.0001 (0.02) | [−0.03, 0.03] | 1.00 [0.97, 1.03] | $\chi^2(1) = 0.00$ | 0.99 |
| Average pair arousal | −0.13 (0.05) | [−0.22, −0.03] | 0.88 [0.80, 0.97] | $\chi^2(1) = 7.08$ | 0.008** |
| Average pair valence × Average pair arousal | −0.007 (0.03) | [−0.07, 0.05] | 0.99 [0.94, 1.06] | $\chi^2(1) = 0.05$ | 0.83 |

*Note.* Temporal distance memory was modeled with linear mixed-effects models (*F* tests); temporal order memory was modeled with logistic mixed-effects models (Wald $\chi^2$ tests). OR = odds ratio. 95% CIs are Wald intervals. **$p < 0.01$.

**Supplementary Table 3**

*Mixed-effects modeling results on temporal memory effects, controlling for valence, arousal, or their interaction*

| Effect and control variable | *B* (*SE*) | 95% CI | OR [95% CI] | Test | *p* |
|---|---|---|---|---|---|
| **Affective spillover ~ Within-sequence temporal memory** | | | | | |
| ***Temporal distance memory*** | | | | | |
| + Sequence valence | −0.01 (0.005) | [−0.02, −0.001] | — | $F = 4.85$ | 0.03* |
| + Sequence arousal | −0.01 (0.005) | [−0.02, −0.001] | — | $F = 4.84$ | 0.03* |
| + Sequence valence × Sequence arousal | −0.01 (0.005) | [−0.02, −0.001] | — | $F = 4.82$ | 0.03* |
| ***Temporal order memory*** | | | | | |
| + Sequence valence | 0.03 (0.02) | [0.001, 0.07] | — | $F = 4.13$ | 0.04* |
| + Sequence arousal | 0.03 (0.02) | [0.002, 0.07] | — | $F = 4.23$ | 0.04* |
| + Sequence valence × Sequence arousal | 0.03 (0.02) | [0.001, 0.07] | — | $F = 4.09$ | 0.04* |
| **Across-sequence temporal memory ~ First- × second-sequence valence** | | | | | |
| ***Temporal distance memory*** | | | | | |
| + Average pair valence | −0.93 (0.07) | [−1.07, −0.80] | — | $F = 184.09$ | < 0.001*** |
| + Average pair arousal | −0.93 (0.07) | [−1.07, −0.80] | — | $F = 183.97$ | < 0.001*** |
| + Average pair valence × Average pair arousal | −0.93 (0.07) | [−1.06, −0.79] | — | $F = 181.64$ | < 0.001*** |
| ***Temporal order memory*** | | | | | |
| + Average pair valence | −0.44 (0.10) | [−0.64, −0.24] | 0.64 [0.53, 0.78] | $\chi^2(1) = 19.32$ | < 0.001*** |
| + Average pair arousal | −0.44 (0.10) | [−0.63, −0.24] | 0.65 [0.53, 0.79] | $\chi^2(1) = 19.13$ | < 0.001*** |
| + Average pair valence × Average pair arousal | −0.44 (0.10) | [−0.64, −0.24] | 0.64 [0.53, 0.78] | $\chi^2(1) = 19.21$ | < 0.001*** |

*Note.* Temporal distance memory was modeled with linear mixed-effects models (*F* tests); temporal order memory was modeled with logistic mixed-effects models (Wald $\chi^2$ tests). OR = odds ratio. 95% CIs are Wald intervals. $^*p < 0.05$; $^{***}p < 0.001$.

Within sequences, semantic similarity was averaged across adjacent image pairs to derive a sequence-level similarity score; across sequences, similarity was computed directly for each image pair. Sequence-level semantic similarity did not predict within-sequence temporal memory (distance memory: $B = -0.48$ ($SE = 0.35$), 95% CI [−1.17, 0.21], $F = 1.87$, $p = 0.17$; order memory: $B = -0.24$ ($SE = 0.49$), 95% CI [−1.19, 0.71], OR = 0.79, 95% CI [0.30, 2.04], $\chi^2(1) = 0.25$, $p = 0.62$). Importantly, controlling for semantic similarity did not alter the associations between temporal memory and affective spillover (distance memory: $B = -0.01$ ($SE = 0.005$), 95% CI [−0.02, −0.001], $F = 4.76$, $p = 0.03$; order memory: $B = 0.04$ ($SE = 0.02$), 95% CI [0.002, 0.07], $F = 4.32$, $p = 0.04$). Across sequences, higher semantic similarity predicted shorter remembered temporal distance between images ($B = -0.39$ ($SE = 0.18$), 95% CI [−0.74, −0.04], $F = 4.85$, $p = 0.03$), replicating prior findings[26]. Critically, however, emotional valence-transition effects on temporal memory remained significant after controlling for semantic similarity (distance memory: $F = 180.98$, $p < 0.001$; order memory: $\chi^2(1) = 19.76$, $p < 0.001$).

*Temporal memory: Item exposure history.* Given that individual emotional images could be probed multiple times during temporal memory testing (see *Methods: Within- vs. across-sequence pairs* in the main text), we examined whether our primary findings on temporal memory remained robust after accounting for item exposure history. We quantified item exposure history as the total number of prior memory-test exposures for the two items in each test pair. We then included it as a control variable when examining 1) event boundary effects in temporal distance/order memory, 2) the associations between temporal distance/order memory and affective spillover, 3) the associations between alpha burst time and temporal distance/order memory, and 4) the effects of emotional-valence shifts on temporal distance/order memory. All the result patterns remained significant ($p$s < 0.05), suggesting these effects were robust to differences in item exposure history. See Supplementary Table 4 below for full model statistics.

*Mixed-effects modeling: Random-effects structure for primary analyses.* Because we used a data-driven approach to determine whether to include by-participant, by-face, or by-sequence random slopes, we report below the full model specifications, including the random-effects structure, for each primary analysis for transparency (see Supplementary Table 5 below).

**Supplementary Table 4**

*Mixed-effects modeling results on temporal memory effects, controlling for item exposure history*

| Outcome and predictor | *B* (*SE*) | 95% CI | OR [95% CI] | Test | *p* |
|---|---|---|---|---|---|
| **Temporal distance memory (Fig. 3b)** | | | | | |
| Boundary (within vs. across) | −0.54 (0.06) | [−0.66, −0.43] | — | $F = 87.24$ | $< 0.001$*** |
| Item exposure history | −0.07 (0.01) | [−0.09, −0.05] | — | $F = 49.58$ | $< 0.001$*** |
| **Temporal order memory (Fig. 3c)** | | | | | |
| Boundary (within vs. across) | 0.78 (0.11) | [0.57, 0.99] | 2.19 [1.78, 2.70] | $\chi^2(1) = 54.03$ | $< 0.001$*** |
| Item exposure history | 0.009 (0.01) | [−0.02, 0.04] | 1.01 [0.98, 1.04] | $\chi^2(1) = 0.45$ | 0.50 |
| **Affective spillover (Fig. 4a)** | | | | | |
| Within-sequence temporal distance memory | −0.010 (0.005) | [−0.020, −0.001] | — | $F = 4.47$ | 0.03* |
| Item exposure history | 0.009 (0.006) | [−0.002, 0.020] | — | $F = 2.54$ | 0.11 |
| **Affective spillover (Fig. 4b)** | | | | | |
| Within-sequence temporal order memory (correct vs. incorrect) | 0.03 (0.02) | [0.003, 0.070] | — | $F = 4.26$ | 0.04* |
| Item exposure history | 0.010 (0.006) | [−0.004, 0.020] | — | $F = 2.91$ | 0.09 |
| **Within-sequence temporal distance memory (Fig. 5c)** | | | | | |
| Alpha burst time | 2.13 (0.34) | [1.46, 2.79] | — | $F = 39.58$ | $< 0.001$*** |
| Item exposure history | −0.07 (0.02) | [−0.10, −0.04] | — | $F = 17.71$ | $< 0.001$*** |
| **Within-sequence temporal order memory (Fig. 5d)** | | | | | |
| Alpha burst time | −2.32 (0.48) | [−3.26, −1.38] | 0.10 [0.04, 0.25] | $\chi^2(1) = 23.46$ | $< 0.001$*** |
| Item exposure history | 0.02 (0.02) | [−0.03, 0.07] | 1.02 [0.97, 1.07] | $\chi^2(1) = 0.71$ | 0.40 |
| **Across-sequence temporal distance memory (Fig. 6a)** | | | | | |
| First- × second-sequence valence | — | — | — | $F = 185.92$ | $< 0.001$*** |
| Valence shift vs. no shift | 0.47 (0.03) | [0.40, 0.53] | — | $t = 13.63$ | $< 0.001$*** |
| Positive-to-negative vs. negative-to-positive | −0.06 (0.05) | [−0.15, 0.03] | — | $t = -1.23$ | 0.22 |
| Item exposure history | −0.07 (0.01) | [−0.09, −0.04] | — | $F = 26.33$ | $< 0.001$*** |
| **Across-sequence temporal order memory (Fig. 6b)** | | | | | |
| First- × second-sequence valence | — | — | — | $\chi^2(1) = 19.32$ | $< 0.001$*** |
| Valence shift vs. no shift | 0.22 (0.05) | [0.12, 0.32] | 1.25 [1.13, 1.37] | $z = 4.40$ | $< 0.001$*** |
| Positive-to-negative vs. negative-to-positive | 0.16 (0.07) | [0.03, 0.29] | 1.17 [1.03, 1.34] | $z = 2.35$ | 0.02* |
| Item exposure history | −0.0007 (0.02) | [−0.04, 0.04] | 1.00 [0.96, 1.04] | $\chi^2(1) = 0.001$ | 0.97 |

*Note:* Temporal distance memory was modeled with linear mixed-effects models ($F$ tests); temporal order memory was modeled with logistic mixed-effects models (Wald $\chi^2$ tests). OR = odds ratio. 95% CIs are Wald intervals. To facilitate cross-referencing, the corresponding figure number for each effect in the main manuscript is included in parentheses. *$p < 0.05$; ***$p < 0.001$.

**Supplementary Table 5**

*Mixed-effects modeling: Random-effects structure for primary analyses*

| |
|---|
| face rating ~ emotional-sequence valence (Fig. 3a)<br>faceRating ~ seqValCat + list + (1 + seqValCat \| subject) + (1 + seqValCat \| face) |
| temporal distance memory ~ boundary (Fig. 3b)<br>distRating ~ boundary + list + (1 + boundary \| subject) |
| temporal order memory ~ boundary (Fig. 3c)<br>orderMem ~ boundary + list + (1 + boundary \| subject) |
| affective spillover ~ within-sequence temporal distance memory (Fig. 4a)<br>spillover ~ distRating_within + list + (1 \| subject) + (1 \| face) |
| affective spillover ~ within-sequence temporal order memory (Fig. 4b)<br>spillover ~ orderMem_within + list + (1 \| subject) + (1 \| face) |
| within-sequence temporal distance memory ~ alpha burst time (Fig. 5c)<br>distRating_within ~ alpha_burst_time + list + (1 \| subject) + (1 \| sequence) |
| within-sequence temporal order memory ~ alpha burst time (Fig. 5d)<br>orderMem_within ~ alpha_burst_time + list + (1 \| subject) + (1 \| sequence) |
| affective spillover ~ alpha burst time × emotional-sequence valence (Fig. 5e)<br>spillover ~ alpha_burst_time*seqValCat + list + (1 \| subject) + (1 \| sequence) |
| across-sequence temporal distance memory ~ first-sequence valence × second-sequence valence (Fig.6a)<br>distRating_across ~ valCat1_across * valCat2_across + list + (1 \| subject) |
| across-sequence temporal order memory ~ first-sequence valence × second-sequence valence (Fig.6b)<br>orderMem_across ~ valCat1_across * valCat2_across + list + (1 \| subject) |

*Note:* To facilitate cross-referencing, the corresponding figure number for each effect in the main manuscript is included in parentheses.
